# Location-Dependent Differences in Cardiac and Skeletal Muscle Dysfunction Associated With Truncating Titin (*ttn.2*) Variants

**DOI:** 10.1101/2024.12.20.629850

**Authors:** Celine F. Santiago, Inken G Huttner, Ailbhe O Brien, Pauline M. Bennett, Jasmina Cvetkovska, Renee Chand, Mark Holt, Gunjan Trivedi, Louis W. Wang, Kelly A. Smith, Mathias Gautel, Diane Fatkin, Yaniv Hinits

## Abstract

**Background:** Truncating variants (*TTN*tv) in the *TTN* gene, encoding the giant sarcomeric protein titin, cause a range of human cardiac and skeletal muscle disorders of varying penetrance and severity. The effects of variant location on clinical manifestations are incompletely understood.

**Methods:** We generated six zebrafish lines carrying tv in the *ttn.2* gene at the Z-disk, I-band, A-band, and M-band titin regions. Expression of titin transcripts was evaluated using qPCR. Phenotype analysis was performed during embryonic development and in adult hearts.

**Results:** Using location-specific primers, we found a significant reduction of Z-disk and I-band *ttn.2* transcripts in all homozygous embryos but levels of A-band and M-band transcripts were reduced only in lines with truncations distal to the *cronos* promoter. Homozygous embryos uniformly died by 7-10 days post fertilization with marked impairment of cardiac morphology and function. Skeletal muscle motility and sarcomere organization were more disrupted in mutants with truncations distal to the *cronos* promoter compared to those proximal. The most C-terminal line (e232), which lacked only the titin kinase and M-band regions, differed from other lines, with homozygous embryos showing incorporation of truncated Ttn.2/Cronos protein and normal sarcomere assembly, but selective degradation of fast skeletal muscle sarcomeres. Heterozygous embryos were phenotypically indistinguishable from wild-type. High-frequency echocardiography in adult heterozygous fish showed reduced ventricular contraction under resting conditions in A-band mutants. Heterozygous Z-disk and I-band mutants had no significant baseline impairment but were unable to augment ventricular contraction in response to acute adrenaline exposure, indicating a lack of cardiac reserve.

**Conclusions:** Our data suggest that cardiac and skeletal muscle dysfunction associated with truncating *ttn.2* variants is influenced by age, variant location, and the amount of functional titin protein. The distinctive phenotype associated with distal C-terminal truncations may reflect different requirement for C-terminal titin for maintenance of fast, slow and cardiac muscle sarcomeres.

## Introduction

Titin is a giant protein that spans half-sarcomeres from the Z-disk to the M-band in the heart and skeletal muscle. It is a key determinant of passive muscle stiffness and active contraction in health and disease ^1^. In recent years, clinical interest in titin has escalated due to discoveries of genetic variants in patients with cardiac and skeletal myopathies. Truncating variants in the *TTN* gene (*TTN*tv) are now known to be the most common genetic risk factor for dilated cardiomyopathy (DCM) ^2, 3^. While *TTN*tv-associated DCM is typically a dominant adult-onset condition, albeit with variable penetrance, individuals who have two abnormal *TTN* alleles leading to truncated proteins (compound heterozygotes or homozygotes) present with childhood-onset skeletal myopathy ^4–8^. *TTN*tv also cause cardiac and skeletal myopathy in compound heterozygosity with destabilising titin missense mutations ^5, 7^. *TTN*tv have been reported in up to 3% of the general population ^2^ and frequently arise as incidental findings in genetic testing for unrelated disorders and many parental carriers of *TTN*tv remain asymptomatic beyond their sixties.

Effects due to variant location have been proposed as a useful way to stratify *TTN*tv. Most DCM-associated *TTN*tv arise in the titin A-band region, while skeletal muscle-associated variants were proposed to cluster in the M-band ^2–4, 9^. This simplistic view does not account for the presence of *TTN*tv in non-A-band locations in DCM patients, nor A-band *TTN*tv in healthy subjects. Although most patients with A-band *TTN*tv have cardiac presentations, there are increasing reports of overt or subclinical skeletal muscle involvement ^10^. Further, children with congenital recessive *TTN*tv-associated skeletal myopathy may progressively develop DCM ^4, 8^. There are numerous developmental and tissue-specific titin isoforms that arise due to complex alternate splicing. The extent to which any single exon is represented across the range of titin isoforms is indicated by the percent spliced-in (PSI) score. DCM-associated *TTN*tv are thought to be preferentially distributed in exons that have high PSI scores in adult heart ^3^. Whether PSI scores are useful in the context of skeletal myopathy has yet to be clarified. Identifying the subset of *TTN*tv that have the greatest risk of pathogenicity is therefore crucial for informed medical management and genetics counselling of patients and their families.

Zebrafish are a useful animal model to study *TTN*tv effects. There are two titin genes in zebrafish, *ttn.2* (*ttna*) and *ttn.1* (*ttnb*). During early development, *ttn.2* is expressed in both cardiac and skeletal muscle and is the primary titin gene in the heart ^11^. Both heart and skeletal muscle phenotypes have been reported in homozygous *ttn.2* mutants ^12^. We have previously reported adult-onset DCM in heterozygous *ttn.2* zebrafish generated to carry a human A-band *TTN*tv ^13^. *ttn.1* is mainly expressed in skeletal muscle and *ttn.1* mutants have been reported to have predominant skeletal myopathy phenotypes ^14, 15^. An alternative promoter in *ttn.2* encodes Cronos, a short titin isoform comprising A-band and M-band domains. First discovered in zebrafish, the Cronos isoform is conserved in mouse and human ^16, 17^. *Cronos* transcripts are abundant in skeletal muscle, but low in developing hearts.

The aim of our study was to investigate the impact of variant location on cardiac and skeletal muscle structure and function in embryonic and adult zebrafish, using a series of mutant lines carrying *ttn.2* variants predicted to truncate Z-disk, I-band, A-band, and M-band regions.

## Methods

Detailed methods are provided in the Supplementary Material. Data are available from the corresponding authors on reasonable request.

## Results

### *ttn.2* zebrafish models

We generated a series of six zebrafish *ttn.2* lines carrying variants predicted to result in a truncated titin protein (Figures 1A, S1). The e5 line has a nonsense mutation that introduces a premature stop codon in exon 5 before the Z-repeats in the proximal Z-disk region. The e25 line has a 2 bp deletion and premature stop codon in exon 25 in the proximal I-band. In the e105 line, a 7 bp deletion in exon 105 is predicted to shift the reading frame with 12 residues of neo-sequence followed by a premature stop codon at the amino acid corresponding to human *TTNtv*, p.Y12304*. Exon 105 is in the distal I-band proximal to the *cronos* promoter. The e129 line has a 5 bp deletion and stop codon in exon 129 distal to the *cronos* promoter near the I-band/A-band junction. The e201 line, modelling the human *TTN*tv p.R26331*, has previously been described ^13^ and carries an 8 bp deletion and stop codon in exon 201 in the mid A-band. The e232 line has a 1 bp deletion and 8 bp insertion in exon 232 in the distal A-band, upstream of the titin kinase domain and M-band ^18^. Zebrafish exons targeted in these mutants have high sequence homology to the corresponding human *TTN* exons, all of which have high PSI scores in adult human heart (Table S5).

**Figure 1.**
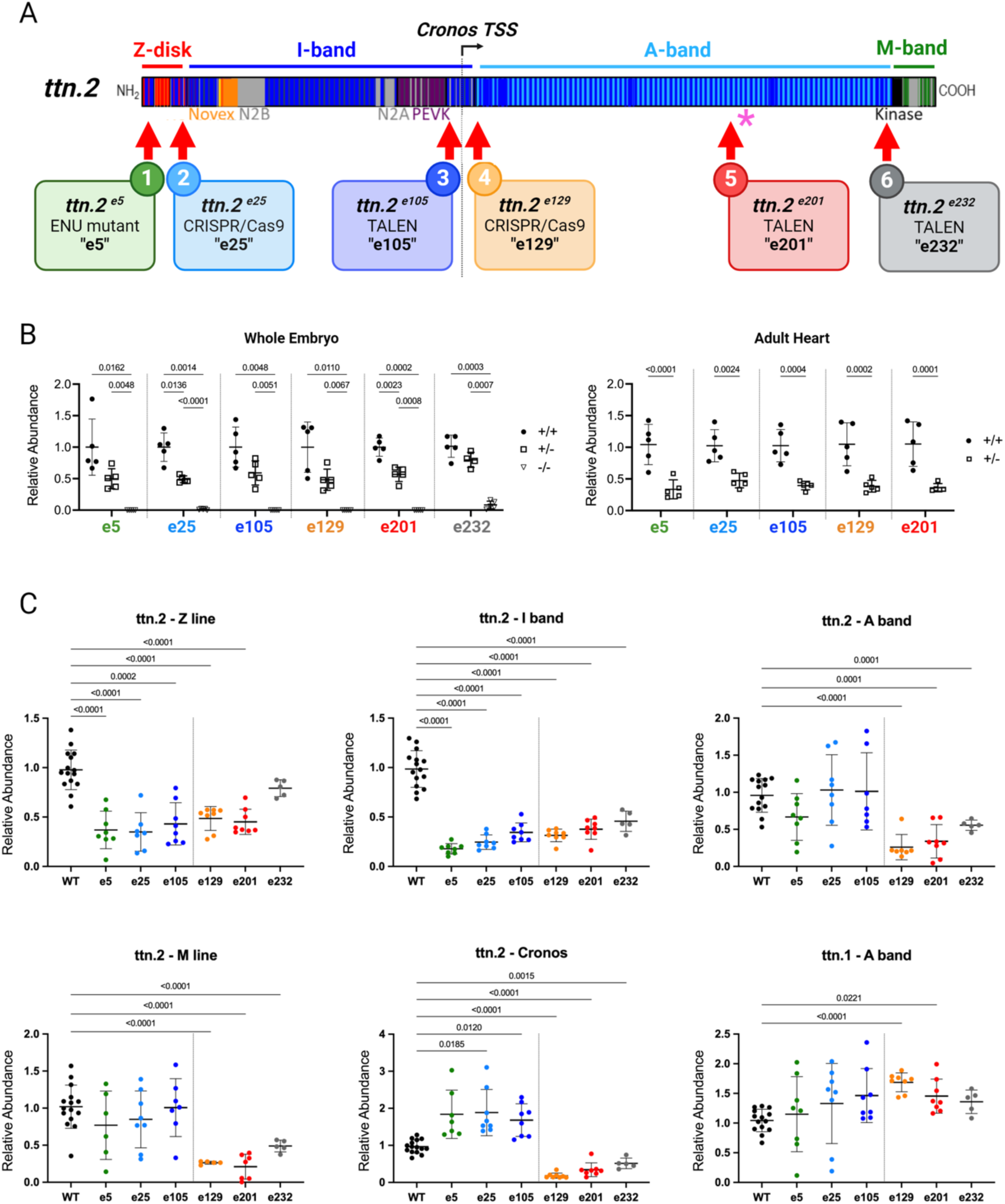
Zebrafish *ttn.2* truncating variants. **A,** Schematic of the Ttn.2 protein showing major domains and the *cronos* translation start site (TSS). Locations of *ttn.2* truncations (red arrows) and the *ttn^x071^* mutant ^14^ (asterisk) are indicated. **B,** qPCR analysis of homozygous (-/-), heterozygous (+/-) and wild-type (WT; +/+) whole embryos (left panel) and adult hearts (right panel) using primers specific for the wild-type allele. **C,** qPCR analysis of homozygous *ttn.2* whole embryos and WT controls using primer pairs specific for various *ttn.2* and *ttn.1* regions. qPCR data (**B,C**) were normalized to house-keeping genes and expressed as absolute levels of abundance. Two-way ANOVA, mean ± SD.

Wild-type (WT) and mutant *ttn.2* alleles were quantified by qPCR analysis of pooled whole embryos (3-5 dpf). Using WT allele-specific primers, levels of WT *ttn.2* transcript were reduced by ∼50% in heterozygous (+/-) mutants and absent in homozygous (-/-) mutants (Figure 1B). In e129-/-, e201-/- and e232 -/- embryos, titin transcripts were globally reduced as indicated by qPCR using location-specific primers across Z-, I-, A- and M-band regions. However, in e5-/-, e25-/- and e105-/- embryos, only Z- and I-band transcripts were reduced, while A- and M-band transcripts were expressed at normal levels (Figure 1C). We hypothesized that these differences might be explained by differential upregulation of the titin Cronos isoform ^16^. Indeed, quantification of *cronos* transcripts revealed significantly increased expression in e5-/-, e25-/- and e105-/- embryos (proximal to the *cronos* promoter) but not in e129-/-, e201-/- and e232-/- embryos (distal to the *cronos* promoter) (Figure 1C). This may suggest up-regulation of *cronos* due to mechanism triggered by degraded mutant mRNA ^19^. There was modest up-regulation of the WT *ttn.1* transcript in all lines but this was statistically significant only for e129-/- and e201-/- (Figure 1C).

### Embryonic cardiac phenotypes

All in-crosses of heterozygous alleles gave rise to Mendelian-ratio progenies. Homozygous embryos from e5, e25, e105, e129 and e201 lines were visually identified from 24 hpf by a dent in the yolk caused by pericardial effusion shortly after the heart tube begins to contract at 23 hpf ^20^. By 3 dpf, there was severe pericardial edema, an indicator of impaired ventricular contraction, as well as small eyes and craniofacial dysplasia (Figure 2A, B). The heart chambers were small and linearly arranged, indicating poor development and impaired looping (Figure 2B). e232-/- embryos were a notable exception, with overall wild-type appearance and cardiac morphology. All homozygous embryos died between 4 to 10 dpf (Figure 2C); e232-/- mutants died as larvae but survived relatively longer than other lines. There were no differences in survival between heterozygotesand WT siblings.

**Figure 2.**
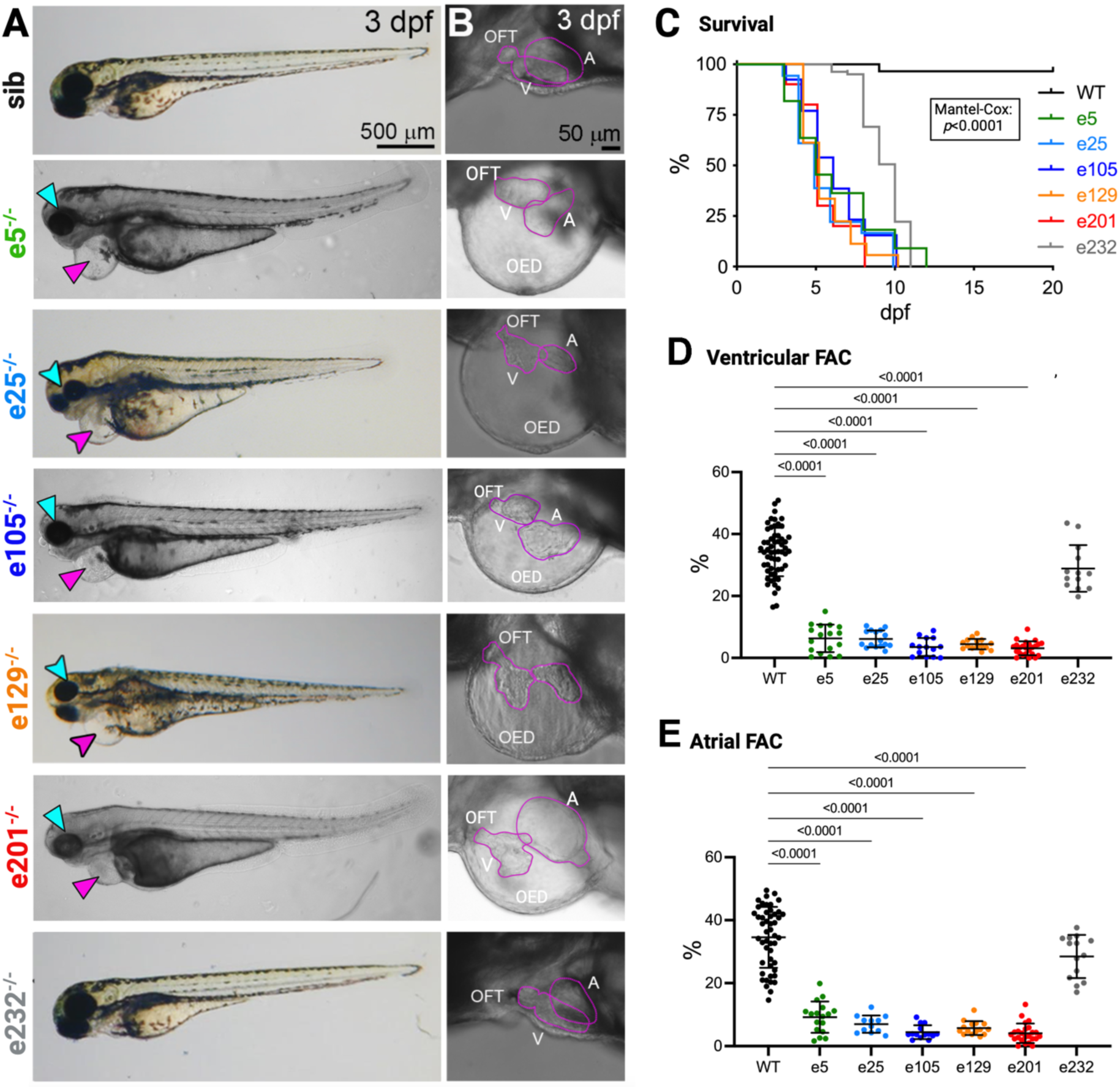
Cardiac morphology and function in homozygous *ttn.2* embryos. **A,** Brightfield images of 3 dpf homozygous and sibling control embryos shown in lateral view, anterior to left, with (**B**) higher magnification of the heart. With the exception of e232-/-, mutant embryos show pericardial edema (magenta arrowhead), as well as small eyes and head (cyan arrowhead). **C,** Survival in homozygous and wild-type (WT) embryos between 0 and 20 dpf; Mantel-Cox test. Ventricular (**D**) and atrial (**E**) contractile function were assessed using video-microscopy. Data are expressed as fractional area change (FAC; %). Unpaired t-tests, mean ± SD. A denotes atrium, OED, edema, OFT, outflow tract, and V, ventricle.

Cardiac function was evaluated in embryonic fish using video-microscopy. For all lines except e232, homozygous mutants showed a marked reduction in ventricular and atrial contractile function (assessed by fractional area change) from 3 dpf onwards (Figure 2D, E). Cardiac function in heterozygous *ttn.2* embryos was indistinguishable from WT siblings (Figure S3). Heart rates were similar in WT and mutant embryos.

### Myocardial architecture in embryonic fish

Embryonic hearts were evaluated using immunofluorescence staining. At 3 dpf, the intensity of myosin heavy chain (MyHC) staining in e5-/-, e25-/-, e105-/-, e129-/-, and e201-/- embryonic hearts was reduced, with numerous aggregates and few very disorganized fibrils (Figures 3A, 3B, S4A,B). Z-disk titin interacts with the cytoskeletal protein, α-actinin ^21^. In e5-/-, e25-/-, e105-/-, e129-/-, and e201-/- embryos, α-actinin staining showed aggregates and poorly organized striations, suggesting that some Z-I brushes may have formed in the absence of titin (Figure 3A, 3B, 3D). Myosin binding protein C (MyBP-C), which is in close contact with the C-zone of A-band titin ^18^, was absent from the cardiac sarcomeres of these mutants (Figure 3C). Intriguingly, this suggests that despite the apparent absence of direct titin-MyBP-C contacts in the assembled (relaxed) thick filaments, A-band titin is indispensable for the correct formation of the myosin-MyBP-C complex *in vivo.* Myomesin-2 (Myom2) interacts with titin in the M-band in normal hearts (reviewed in ^22^). Myom2 was disorganized and present only at very low levels, suggestive of impaired M-band myosin-titin arrays (Figure 3D).

**Figure 3.**
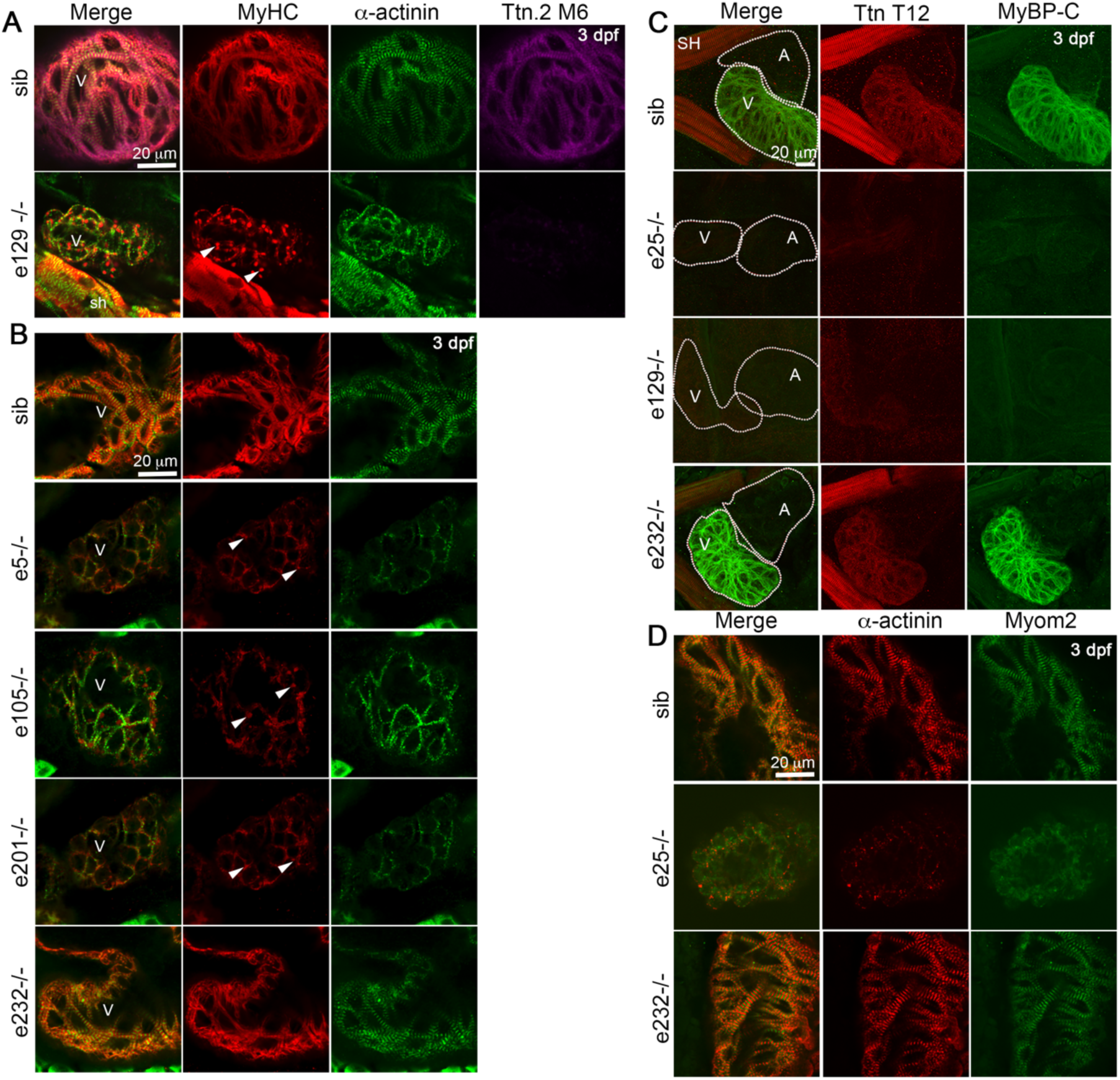
Cardiac sarcomere assembly in *ttn.2* embryos. **A** and **B,** Single confocal scans through ventricular sarcomeres of 3 dpf homozygous and sibling (sib) control embryos, immunostained for myosin heavy chain (MyHC), α-actinin, and Ttn.2 M6. Mutants show punctate aggregates (arrowheads) except e232-/- embryos in which sarcomeres appear normal. No titin protein is detected by Ttn.2 M6 antibodies in e129-/- hearts. **C,** Confocal stacks of 3 dpf hearts of homozygous *ttn.2* and sibling control embryos, immunostained for Ttn T12 and myosin binding protein-C (MyBP-C), shown in ventral view, anterior to left. Heart chambers show very low levels of Ttn T12 and MyBP-C. **D**, Single confocal scans through ventricular sarcomeres of e25-/-, e232-/-, and sibling control embryos, immunostained for α-actinin and myomesin-2 (Myom2). e25-/- embryos show punctate aggregates of α-actinin and only low levels of Myom2. In (**C**) and (**D**), cardiac morphology and sarcomeres in e232-/- embryos are similar to control. A denotes atrium, SH, sternohyoideus muscle, and V, ventricle.

Cardiac expression of Ttn.2, Ttn.1, and Cronos was evaluated using *in situ* hybridization and immunostaining. In WT embryos, there was strong staining using the T12 antibody recognizing an epitope in I2-I3 at the start of the I-band in both Ttn.2 and Ttn.1 ^23^ (Figure 3C), and a novel antibody that recognizes the Ttn.2 M6 M-band domain (Figures 3A, S5A,B). Another antibody against the Ttn.1 M8-M9 region, but detecting both Ttn.1 and Ttn.2 (Figures S2, S5A,C), showed normal cardiac staining but reduced M-band staining in *ttn.1* e7-/- embryos somitic muscle sarcomeres that express both Ttn.1 and Ttn.2 (Figures S2B, S6M-X). In WT embryos, *ttn.1* transcripts express only weakly in the heart at 1 dpf (Figure S6B, S6D, S6F and ^11^) with negligible expression by 3 dpf (Figure S6H, S6J, S6L). Like previous studies ^14^, we found that embryonic heart development in *ttn.1* mutant lines is normal (Figure S2C, C’). These finding suggest Ttn.2 but not Ttn.1 is essential for normal sarcomere assembly and formation of a functional heart.

For all homozygous embryos except e232-/-, there was no detectable Ttn.2 protein or Ttn.1 in the heart above background levels (Figure 3A, 3C and S5). Our qPCR data in whole embryos (Figure 1) suggested that *cronos* upregulation might be a potential compensatory mechanism for titin deficiency. However, these changes appear to be driven primarily by *cronos* expression in skeletal muscle, since in situ hybridization failed to detect *cronos* in the heart (Figure S6Y, S6Z and ^16^).

In keeping with preserved cardiac function, sarcomere organization was indistinguishable in T12 stained e232-/- embryos compared to WT siblings (Figure 3B). Given the negligible expression of Ttn.1 and Cronos in 3 dpf hearts, we speculate that e232 transcripts may be incompletely degraded and that truncated e232 Ttn.2 protein is incorporated into cardiac sarcomeres and able to facilitate near-normal organization of myosin and sarcomeric proteins such as Myom2 (Figure 3D).

### Skeletal muscle embryonic phenotypes

Skeletal muscle motility and structure was evaluated in *ttn.2* embryos. Embryos with truncating variants proximal to the *cronos* promoter (e5-/-, e25-/-, e105-/-) had reduced motility compared with sibling control embryos at ∼24hpf, which was limited to just a tremor when stimulated by touch by 3 dpf (Figure S7A). Immunofluorescence staining for myosin and actin in these 3 dpf mutants showed structured sarcomeres, but some myofibrils were mis-aligned and break points were evident in the middle of both slow and fast muscle fibers (Figures 4A, S7B-D). MyBP-C and α-actinin appeared normal (Figure S7E, S7F). Immunostaining with T12, recognizing an epitope 3’ of the *ttn.2* truncation site, showed strong staining in both slow and fast muscle fibers, which most likely results from Ttn.1 (Figures S7E and S6N, S6R, S6T, S6X). Ttn.2 M6 was also detected strongly in e5-/-, e25-/- and e105-/- embryos (Figure S7F-H) and is likely to result from Cronos protein. Other M-band proteins localized correctly to the M-band; Myom2 in fast muscle (Figure 4B) and Myom1 in both fast and slow muscle fibers (Figure S7I-K). Overall, these data suggest that Ttn.1 and Cronos have a ruler function and allow many proteins to find their position in the sarcomere in the absence of full-length Ttn.2. However, reduced motility and fiber breakage indicate that sarcomere structure is weaker and prone to damage once contractions commence.

**Figure 4.**
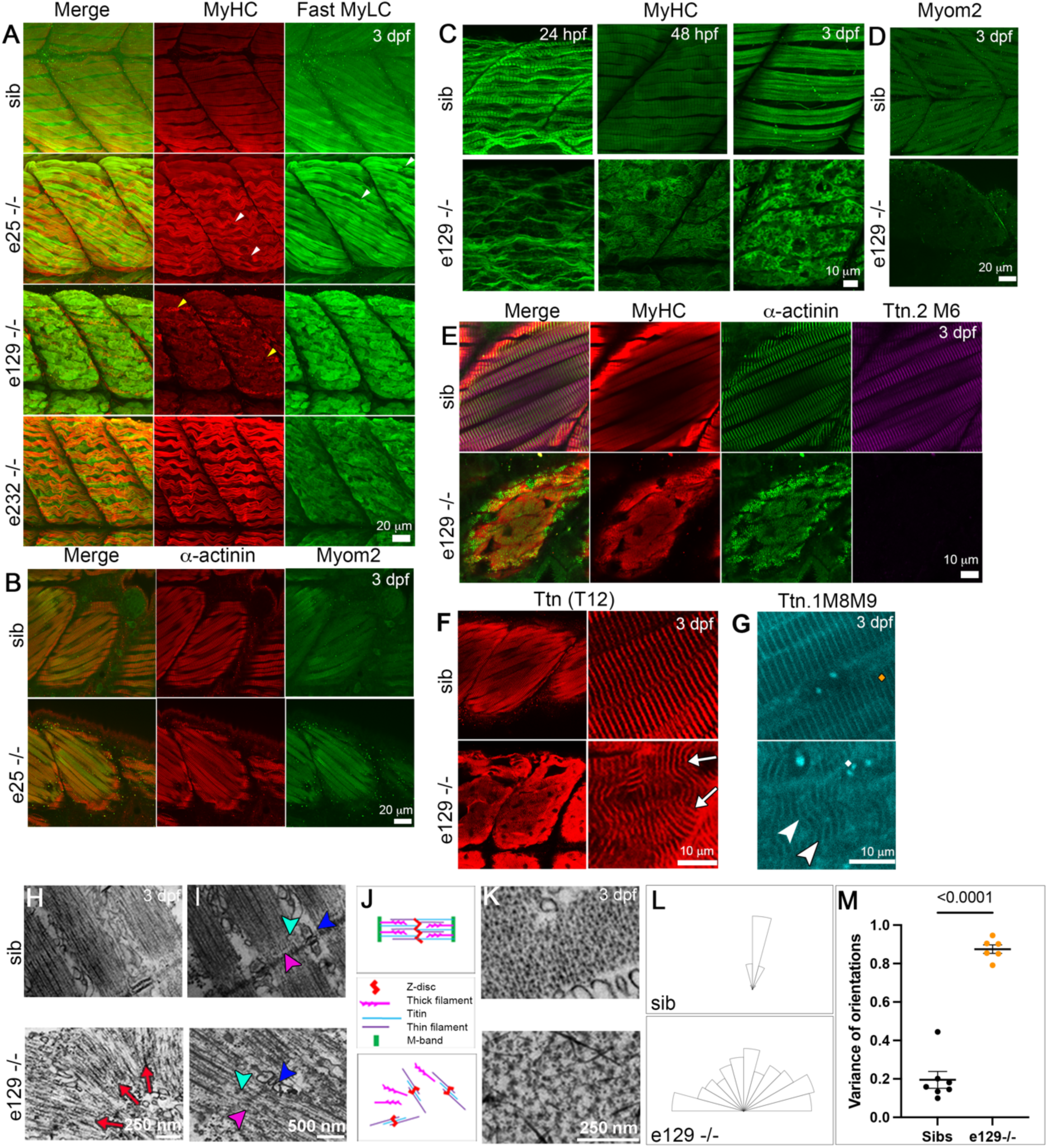
Skeletal muscle phenotypes in *ttn.2* embryos. **A** through **G,** Immunofluorescence staining of 3 dpf homozygous and sibling (sib) control embryos for myosin heavy chain (MyHC), fast myosin light chain (Fast MyLC), α-actinin, myomesin-2 (Myom2), Ttn.2 M6, Ttn T12, and Ttn.1 M8-M9, shown in lateral view, anterior to left. MyHC is strongly expressed in superficial slow fibers, while MyLC labels fast fibers. **A**, e25-/- embryos show some good sarcomeric structure as well as regions of damaged sarcomeres in the middle of the fiber (white arrowheads). e129-/- embryos have disorganized sarcomeres in both slow (yellow arrowheads) and fast fibers, while e232-/- embryos have striated slow fibers but no organized thick filaments in fast fibers. **B**, In e25-/- embryos, α-actinin (slow and fast fibers) and Myom2 (fast only) seem striated. **C** through **G**, In e129-/- embryos, abnormal striation is detected from 24 hpf to 3 dpf; MyHC and α-actinin and show no normal striation but some radial organization is seen around random points in the fiber (**C,E**), Myom2 is abolished (**D**), and Ttn.2 M6 is completely missing (**E**). This abnormal striation is also seen with Ttn T12 (**F**) and Ttn.1 M8-M9 (**G**). **H** through **M,** Electron micrographs of longitudinal sections of 3 dpf e129-/- and sibling control embryos. Thin filaments in mutant embryos are arranged in I-Z-I brushes that are organized in radiating patterns (**H**, red arrows), highlighting structural features (**I**), including triads (blue arrowheads), Z-disks (pink arrowheads) and thin filaments (cyan arrowheads). **J**, Schematic illustrating arrangement of thick and thin filaments. **K,** Electron micrographs of transverse sections of fibers in 3 dpf embryos showing lattice pattern of thick and thin filaments in sibling controls and increased interfilament space in e129-/-. **L** and **M,** Quantification of directionality of fibres. Rose diagram of representative somitic muscle (**L**) and graph comparing the variance of the orientations (**M**). The possible range is 0 to 1, where 0 is completely ordered and 1 is completely random. Unpaired t-tests, mean ± SEM.

### Severe loss of skeletal muscle sarcomeric structure in embryos with *ttn.2* truncations distal to the *cronos* promoter

The severity of skeletal muscle phenotypes was greater in embryos with truncating variants distal to the *cronos* promoter (e129-/-, 201-/-) compared to those with proximal truncations (e5-/-, e25-/-, e105-/-). The former embryos were barely motile at 24 hpf, showing only weak twitches. By 3 dpf, embryos were paralyzed with no response to mechano-stimulation (Figure S8A). Already at 24 hpf, sarcomeres in e129-/- embryos had immature thin myofibrils resembling early differentiated myofibrils ^24^ with only small areas of low-level sarcomeric organization (Figures 4C, S8D). By 2 and 3 dpf, immunofluorescence staining for myosin and actin showed a loss of axial fibrillar organization in both slow and fast fibers (Figures 4A, 4C, and S8B, S8C). Rather, short lengths of ∼1 µm structures were revealed by staining for α-actinin (Figure 4E) and antibodies that detect Ttn.1, such as Z1Z2, T12, and Ttn.1 M8-M9 (Figures 4F, 4G, S8B). The directionality of these structures was random, differing greatly from normal organized sarcomeres (Figure 4L, 4M) and resembling I-Z-I complexes. I-Z-I complexes are structures formed during myofibrillogenesis that contain Z-disk components as well as α-actin and titin. These can be put in register on actin filamentous structures and can form in the absence of MyHC ^25^. Ttn.2 (M6 antibody) was not detected in these mutants (Figures 4E, S8E) while other M-band proteins such as Myom2 were only weakly detected (Figure 4D). This suggests that fibrils expressing only Ttn.1 have lost their rigidity or their ability to associate side by side, so that they grow in a wavering, random way when viewed longitudinally. Electron microscopy supports this, with a few repeats of sarcomere-like structures running in different directions (Figure 4H-4K). Between the Z-disks and their associated thin filaments, groups of thick filaments were seen. However, they did not appear to be organized as A-bands and no M-band structures were seen. Furthermore, the separation of the thick filaments in cross section was greater than seen in WT siblings. The sarcoplasmic reticulum was present, and some triad structures remain associated with Z-disks (Figure 4J). Overall, the results show that in skeletal muscles of homozygous embryos with *ttn.2* truncations distal to the *cronos* promoter, structures containing I-Z-I brushes and Ttn.1 are formed, but Ttn.1 alone cannot support proper organization of A- and M-bands.

### Sarcomere structure disintegrates in fast fibers of embryos with M-band ttn.2 truncation

Similar to the cardiac findings, the skeletal muscle phenotype in e232-/- embryos was distinct to the other mutants. The overall appearance of e232-/- embryos was indistinguishable from WT siblings at 3 dpf (Figure 2A). However, whereas coiling movement within the chorion was normal at 24 hpf, e232-/- embryos had an abnormal swimming pattern at 48 hpf, which involved slower tail movement and less motion of the trunk (Supplementary Video). By 5 dpf, e232-/- embryos lacked burst-swimming upon touch, unlike WT siblings. However, spasms of the trunk, pectoral fin movement and increased eye movement in response to touch were detected, indicating intact sensory reflexes (Figure S9A-S9C). Burst-swimming behaviour is mediated primarily by fast fibers, whilst the early coiling movement is mediated by early differentiating slow muscle ^26, 27^. By 5 dpf, e232-/- mutants developed curvature of the trunk and failed to inflate their swim bladders (Figure 5A, Supplementary Video) ^27^.

**Figure 5.**
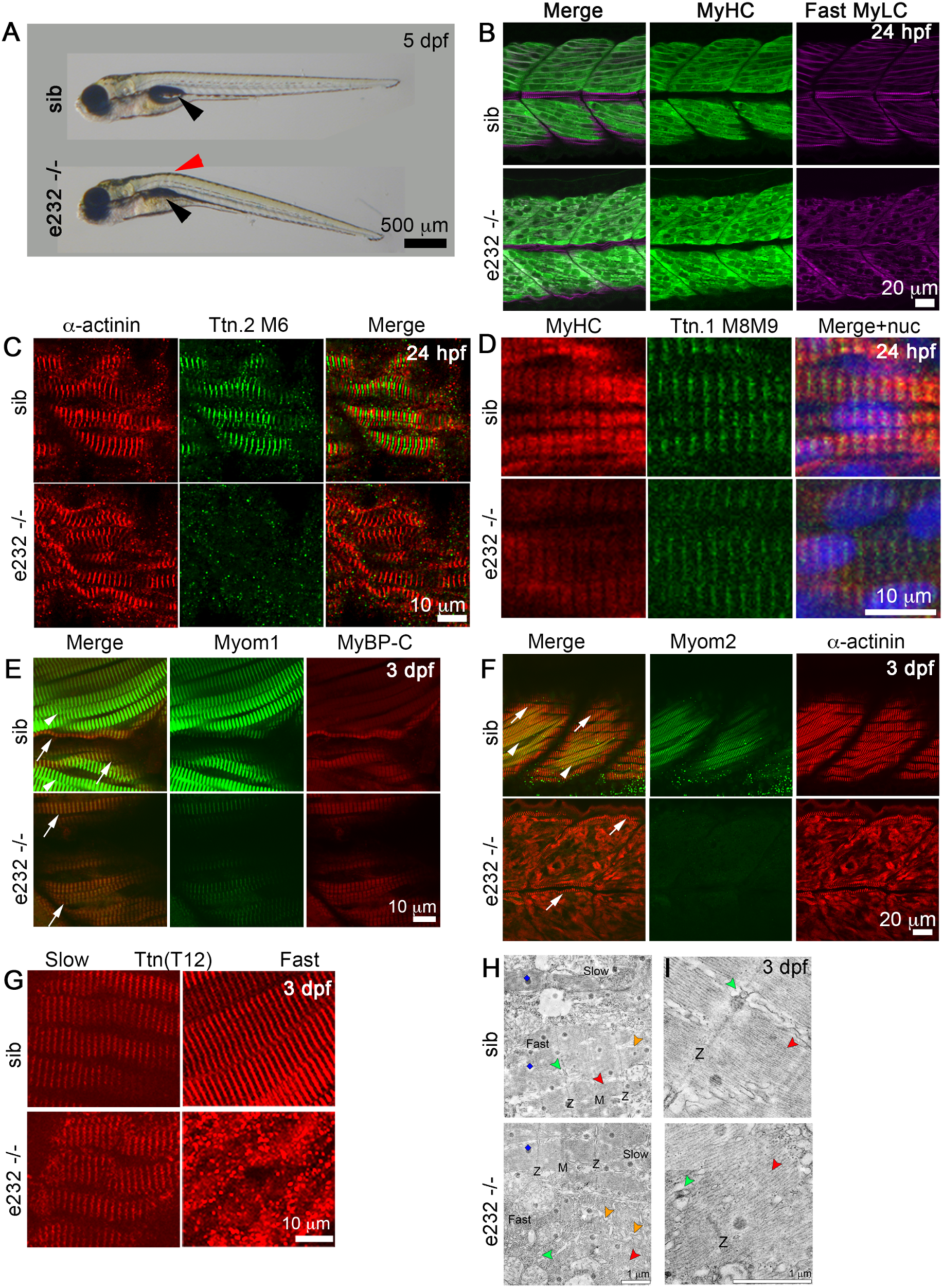
Skeletal muscle phenotypes in e232-/- embryos. **A,** Brightfield images of 5 dpf e232-/- and sibling (sib) control embryos, lateral view, anterior to left. Mutants show a distinctive trunk curvature (red arrowhead) and fail to inflate swim bladder (black arrowheads). **B** through **G,** Immunofluorescence staining of e232-/- and sibling (sib) control embryos for myosin heavy chain (MyHC), fast myosin light chain (Fast MyLC), α-actinin, Ttn.2 M6, Ttn.1 M8-M9, myomesin-1 (Myom1), myosin binding protein-C (MyBP-C), myomesin-2 (Myom2) and Ttn T12, shown in lateral view, anterior to left. At 24 hpf (**B-D**), MyHC, fast MyLC, α-actinin and Ttn.1 M8-M9 (slow) show regular sarcomeres although Ttn.2 M6 is not detected. By 3 dpf (**E-H**), Myom1 (strong in fast fibers) and MyBP-C (strong in slow fibers) are present in both slow (**E**, arrows) and fast fibers (**E**, arrowheads) in controls but are detected only in slow fibers in e232-/- embryos. Myom2 is expressed only in fast fibers (**F**, arrowheads), whereas α-actinin is in both slow (**F**, arrows) and fast (**F**, arrowheads) fibers. In e232-/- embryos, α-actinin is normal in slow fibers (arrows) but is disorganized and aggregated in the deeper fast fibers. Myom2 shows diminished signal above background. **G**, Fibers are scanned at different levels to highlight superficial slow muscle (left panels) and deeper medial fast fibers (right panels). Ttn T12 striation patterns are similar in e232-/- and control embryos in slow fibers but mutants have a punctate disorganised staining fast fibers. **H** through **I,** Electron micrographs of longitudinal muscle sections of 3 dpf e232-/- and sibling control embryos. Control embryos have organized sarcomeric structure with clear Z-disk (Z) and M-band (M) visible in both slow and fast muscle fibers. e232-/- embryos have organized sarcomeres only in slow muscle fibers. Thick filaments (red arrowheads) are present in mutant fast fibers but are not arranged into sarcomeres. Sarcoplasmic reticulum (orange arrowheads) is disordered in mutant fast fibers. Blue kites indicate EM artefacts. Triads (green arrowheads) are located at the Z-disk in control fast fibers. In mutant fast fibers, there are a small number of triads and Z-disk but these are not associated together.

At 24 hpf shortly after differentiation, sarcomeric structure in both slow and fast fibers in e232-/- embryos appeared normal (Figure 5B-5D). As in cardiac sarcomeres, T12 signal was present (Figure 5G) but Ttn.2 M6 was absent (Figure 5C), suggesting that truncated Ttn.2 and Cronos were incorporated into the sarcomere. By 3 dpf, however, while slow muscle fibers maintained their regular sarcomeric structure, fast muscle sarcomeres were severely disrupted as indicated by myosin, MyBP-C, α-actinin and T12 (detecting Ttn.2 upstream from the truncation domain) staining patterns (Figures 4A, 5E-5G). The M-band protein Myom1 is normally present in both slow and fast fibers but was only detected in e232-/- embryo slow fibers (Figure 5E). Myom2, normally present only in fast fibers, was severely diminished in e232-/- embryos (Figure 5F). Electron microscopy revealed normal sarcomeric structure in e232-/- slow muscle fibers, whereas no sarcomeres were visible in fast muscle fibers and thick filaments appeared to be highly disordered (Figure 5H, 5I). A small number of Z-discs and triads were also visible, but not associated.

### Adult cardiac phenotypes

Heterozygous *ttn.2* mutant fish survived to adulthood with overall normal appearance and swimming behaviour. *ttn.2* transcript expression in mutant lines was reduced by ∼50% relative to WT (Figure 1B). Confirming previous studies ^14^, neither reduced levels of titin protein nor truncated titin proteins were detectable using Western blots with an N-terminal specific titin antibody, or Coomassie stained polyacrylamide gels (Figure S10). Due to the lack of embryonic cardiac phenotype in e232-/- fish, e232+/- fish were not included in the adult analyses.

Cardiac function in 12-month-old fish was evaluated using high frequency echocardiography (Figure 6, Table S6). Ventricular size (EDV/BSA) in the heterozygous *ttn.2* fish was unchanged compared to WT, except for e201+/- that showed modest dilatation (Figure 6A), in line with our previous findings ^13^. Significant impairment of ventricular contractile function was seen only for e129+/- and e201+/- lines (Figure 6B through E). In addition to systolic impairment, these lines also had prolonged isovolumic relaxation time, indicating diastolic dysfunction (Figure 6F). Ventricular function in the A-band mutant, e201+/- was similar in *ttn^xu071^* double mutant (*ttn.2* A-band + *ttn.1* Z-disk truncations, ^14^), implicating predominant *ttn.2* effects (Figure S11).

**Figure 6.**
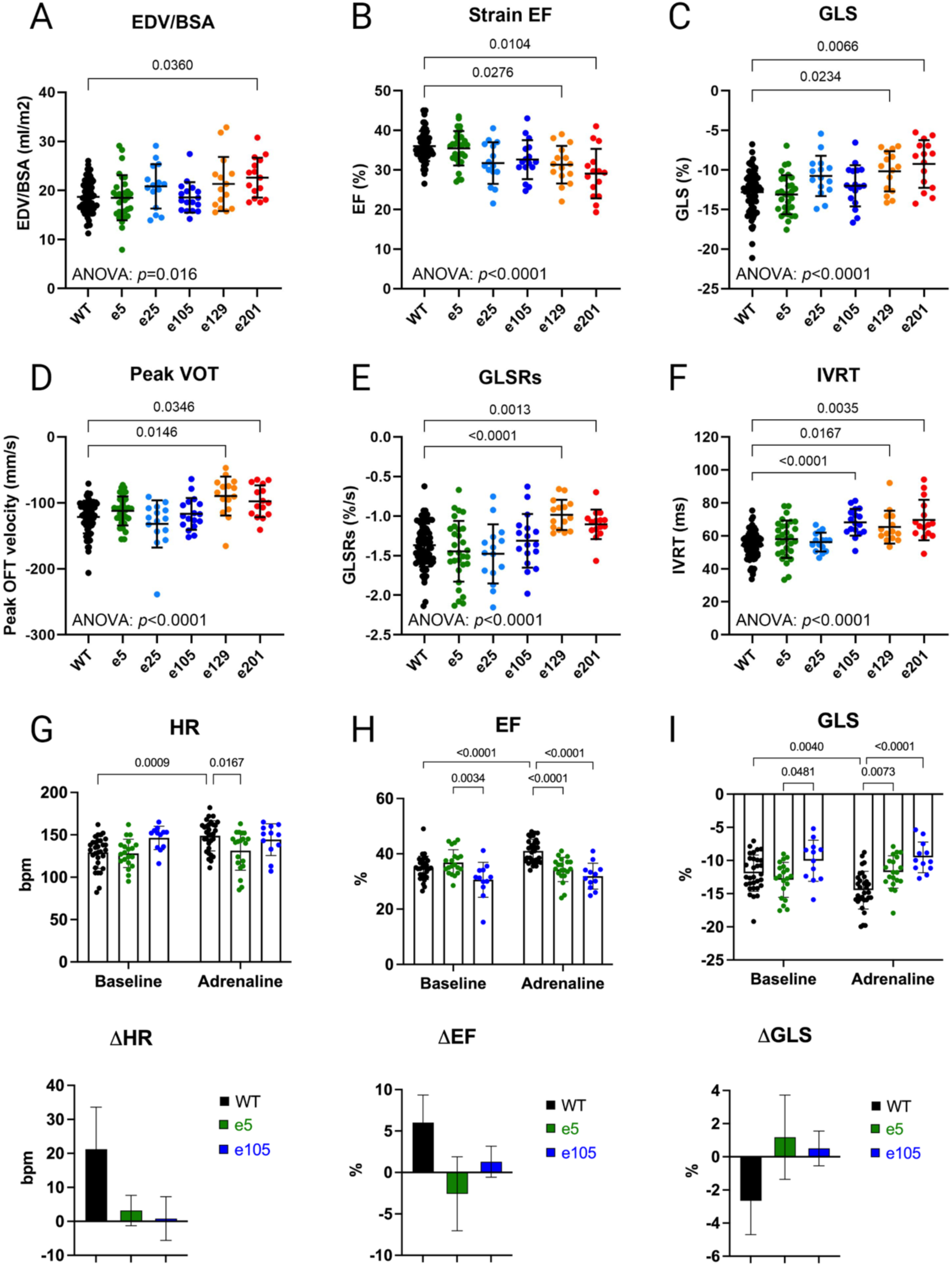
Ventricular size and function in adult heterozygous *ttn.2* zebrafish. Cardiac function was assessed in 12 month-old wild-type (WT) and heterozygous *ttn.2* fish using high frequency echocardiography. **A** through **F s**how ventricular function under resting conditions: **A**, indexed ventricular end-diastolic volume (EDV/BSA), **B**, ejection fraction (EF); **C**, global longitudinal strain (GLS), **D**, peak ventricular outflow tract velocity (VOT), **E**, peak systolic global longitudinal strain rate (GLSRs), **F**, isovolumic relaxation time (IVRT). **G** through **I** show responses (at 2 hr) to acute adrenaline administration in 10-15 month-old fish. Two-way ANOVA, mean ± SD, or Δmean ± SD.

We hypothesized that fish with *ttn.2* truncations outside the A-band might be in a compensated state at baseline, and that exposure to additional stress might unmask latent cardiac dysfunction. We therefore subjected 10-15 month-old e5+/- and e105+/- fish to acute adrenaline stress, as previously described ^28^. WT fish demonstrated the expected adrenaline-induced increases in heart rate and contractility (Figure 6D-F). In contrast, these responses were not seen in e5+/- and e105+/- fish (Figure 6D-F), indicating a lack of cardiac reserve.

Myocardial structure was further interrogated in e5+/- and e201+/- fish, hypothesizing that these two lines are representative of compensated *vs.* penetrant disease states, respectively. No overt differences in wall thickness or trabeculation were observed in haematoxylin and eosin-stained tissue sections in either of these mutant lines relative to WT siblings (Figure S12A, S12D). Sarcomere structure, content and length were also similar to WT (Figure S12B-E).

## Discussion

Zebrafish *TTN*tv models exhibit a varying extent of cardiac and skeletal muscle dysfunction that differs with age, variant location, and the expected amount of normally functioning titin protein. Previous zebrafish studies have focussed mainly on the developmental impact of titin deficiency using homozygous embryos with selected Z-disk and/or A-band *ttn.2* truncations ^12^. We now extend these findings by directly comparing *ttn.2* truncations distributed throughout the titin protein, including the first zebrafish model that truncates the M-band. We also provide new data about cardiac phenotypes in a range of adult heterozygous fish. Our data expand understanding of the roles of titin in heart and skeletal muscle and have potential implications for human titinopathies.

Homozygous *ttn.2* embryos showed marked cardiac dysfunction and early death, similar to reported *ttn.2* mutants ^12^. These findings are most readily explained by a lack of functional Ttn.2 protein in the heart and the lack of effective compensation by Ttn.1 or Cronos. Homozygous *ttn.2* embryos also showed abnormalities of skeletal muscle structure and function, the severity of which appeared to be determined by location with respect to the *cronos* promoter. For proximal truncations, the presence of intact Cronos and Ttn.1 allows formation of fragile sarcomeres that fail following onset of contraction. However, for distal truncations, both Cronos and Ttn.2 are affected and although intact, Ttn.1 appears to be insufficient to support viable sarcomere assembly.

The e232-/- embryos were a notable exception with respect to both the cardiac and skeletal muscle phenotypes. A near-complete A-band in e232-/-, capable of interacting with myosin and organizing MyBP-C, seems essential for incorporation and retention of truncated Ttn.2 in sarcomeres. Remarkably, although loss of full-length Ttn.2 protein prevents sarcomere formation, C-terminal titin appears to be dispensable for cardiomyocyte sarcomere assembly, contraction, and normal heart morphology during early development. In contrast to other lines, minimally truncated Ttn.2 protein is stable enough to allow integration into the sarcomere. Factors such as the truncation position at the distal end of the giant titin protein and the large exon size may result in reduced efficiency of nonsense mediated decay of e232 alleles ^29^. While this might be sufficient to support sarcomere development, the presence of truncated titin seems to affect protein interactions in this region. Homozygous C-terminal titin truncation in mouse develop a late cardiac phenotype and differ greatly from other truncations ^30, 31^. Of note, sarcomeric integration of C-terminal truncated titin has been reported in human muscle biopsies from patients with M-band *TTN*tv associated with autosomal recessive skeletal ± cardiac myopathy ^4–7^.

In e232-/- embryos, sarcomere organization was preserved in cardiac and slow muscle, but not in fast muscle fibers. Since *ttn.2* and *ttn.1* are expressed in both slow and fast fibers, these observations suggest that there are fundamental differences in the requirement for C-terminal titin for the assembly and maintenance in these muscle types. One difference between slow and fast skeletal muscle fibers in the M-band region is expression of myomesin. Whereas Myom1 is present in cardiac, slow and fast muscle in zebrafish, Myom2 is only expressed in fast skeletal muscle, similar to mammalian muscle (reviewed in ^22^). In contrast, Myom2 is present in the murine heart from e14.5 ^22^, while in fish hearts, expression of *myom2a*, one of the fish *MYOM2* homologues, is detected from the early embryonic heart tube stage (Fig. S5AA). Collectively, our data suggest that Myom2 is not essential for sarcomere assembly but may have a role in maintenance of sarcomere organization in fast skeletal muscle. Other proteins that interact with M-band titin, such as obscurin, calpain-3 and myospryn, may be important for differentiating normally functioning sarcomeres in fast versus slow and cardiac muscle fibers. Human patients with C-terminal truncations localize Myom1 and obscurin/Obsl1 to the M-band, but with some disruptions ^7, 32^. Similarly, e232-/- mutants recruited Myom1 to slow and cardiac M-band sarcomeres, possibly via dimerization of Myom1 and binding of Myom1 to obscurin/Obsl1 and myosin. However, absent M-band/titin connections likely gradually result in structural instability and may underpin the disruption seen in human patient samples ^32^. The selective loss of fast skeletal muscle fibers may reflect susceptibility to damage in response to mechanical and metabolic stress ^33^. Interestingly, dystrophin-deficient mdx mice have disproportionate exercise-induced damage in fast skeletal myofibers, and human patients with Salih myopathy and multi minicore disease associated with C-terminal *TTN*tv show a predominance of slow (type I) fibers from a very early age ^4, 5^.

The location of *TTN*tv has emerged as an important issue in human titinopathies. Congenital skeletal titinopathy is recessive and typically associated with biallelic *TTN*tv, except for a rare dominant *TTN*tv in exon 327 ^34^. Cohort studies have demonstrated that these disease-causing *TTN*tv can arise throughout the entire *TTN* gene ^8, 35^. Cardiac involvement is seen in approximately 50% patients and is more likely when both *TTN*tv are in exons that are constitutively expressed in adult heart ^8^. Notably, the age of disease onset is much later in patients in whom at least one of the *TTN*tv is present in the last three M-band exons ^35^. Two autosomal dominant forms of skeletal myopathy, tibial muscular dystrophy and hereditary myopathy with early respiratory failure, have been associated with single *TTN*tv in the final six M-band exons ^8^. These disorders typically occur in adult life and cardiac involvement is variable. Taken together, these findings suggest that *TTN* location and titin “dose” are important determinants of clinical manifestations and disease severity.

The C-terminal part of titin shows unexpected complexity in its integration into the myosin filament, as revealed by recent cryo-electron tomographic (cryoET) structures. Three double strands of titin (dubbed titin alpha and beta) with six molecules in total run along the thick filament ^18^. Both titin alpha and beta interact outside the M-band at the first levels of myosin motor domains. However, whereas titin alpha continues into the M-band, the C-terminal regions of titin beta from about the second Ig-domains after the kinase domain could not be traced in the cryoET structure ^18^, suggesting that titin beta is either truncated or follows a disordered path. The non-equivalence of 3 of the 6 titin filaments of each half thick filament may account for the different impact of truncations N-and C-terminal to the kinase domain. Structural studies of the corresponding zebrafish mutants may therefore provide valuable insight into the different pathomechanism of these mutations.

*TTN*tv are the most common genetic cause of autosomal dominant adult-onset DCM, with the majority of these being in the titin A-band or I/A junction ^2, 3^. Interpretation of the clinical significance of *TTN*tv outside this region has remained a clinical conundrum. Our evaluation of adult zebrafish hearts confirms the importance of A-band *TTN*tv, with significant changes in ventricular contraction seen under resting conditions in e129-/- and e201+/- fish. However, e5+/- and e105+/- fish cannot be considered to have normal hearts as their failure to augment ventricular contraction in response to an adrenaline challenge is indicative of impaired contractile reserve. A recent study found that zebrafish from another *ttn.2* e5+/- mutant had no overt DCM but some mild abnormalities ^36^. All our zebrafish lines had *ttn.2* truncating variants in zebrafish exons that show high sequence homology to constitutively expressed human *TTN* exons. Collectively, our data suggest that *TTN*tv in high PSI cardiac exons confer a genetic susceptibility to DCM and that overt clinical manifestation may depend on patient-related factors such as age, sex, and clinical risk factors, particularly those that increase cardiac mechanical stress. Further human *TTN*tv cohort studies are needed to investigate these points.

Whereas our data, and those from human C-terminal *TTN*tvs ^4–7^, show clear evidence for integration of truncated titin protein into the sarcomere, we have no evidence to support such phenomena for more N-terminal truncations. Recent studies have reported the persistence of some truncated titin protein from DCM *TTN*tv cardiomyocytes, but with conflicting evidence regarding its localization in the sarcomere or in intracellular aggregates ^37–40^. Our study is not without limitations: there remains a lack of definitive evidence for reduced titin protein expression, and for established data for titin exon usage in fetal heart and in fetal and adult skeletal muscle. Due to potential species differences, our findings may not be directly translatable to humans. Nevertheless, our findings provide new insights into the variability underpinning *TTN*tv effects and their potential fundamental impact on thick-filament assembly and are hypothesis-generating for human clinical studies.

## Acknowledgments

We thank Dan Hesselson and Xiaolei Xu for assistance with zebrafish lines; Alexander Alexandrovich and Atsushi Fukuzawa for assistance with antigen expression and purification; Wolfgang Linke for expert assistance with titin protein analysis; the Centre for Ultrastructural Imaging, King’s College London, for electron microscopy services; and the Victor Chang Cardiac Research Institute aquarium staff, in particular Aaron Hay and Cecelia Jenkin, for assistance with zebrafish husbandry. The authors acknowledge the Victor Chang Cardiac Research Institute Innovation Centre, funded by the NSW Government, including the Preclinical Imaging Facility.

