## Supplementary Material File for "Location-Dependent Differences in Cardiac and Skeletal Muscle Dysfunction Associated With Truncating Titin (*ttn.2*) Variants"

TABLE OF CONTENTS


| | | |
|---|---|---|
| Detailed Methods | | 2 |
| Supplemental Figures | | |
| Figure S1 | Generation of zebrafish *ttn.2* truncation variants. | 9 |
| Figure S2 | Zebrafish *ttn.1* truncation variant. | 10 |
| Figure S3 | Cardiac function in heterozygous *ttn.2* mutant embryos. | 11 |
| Figure S4 | Abnormal heart development in *ttn.2* mutants. | 12 |
| Figure S5 | Differences between the Ttn.2 and Ttn.1 M-band regions and specific antibodies. | 13 |
| Figure S6 | Expression of titin in embryonic somitic muscle and heart. | 14 |
| Figure S7 | Homozygous embryos with *ttn.2* tv proximal to the *cronos* promoter have abnormalities in muscle fiber structure and motility. | 16 |
| Figure S8 | Homozygous embryos with *ttn.2* tv distal to the *cronos* promoter have severe abnormalities in muscle structure and function. | 18 |
| Figure S9 | *ttn.2* e232-/- embryos maintain motility of various muscles. | 19 |
| Figure S10 | Evaluation of titin levels in adult zebrafish heart. | 20 |
| Figure S11 | Comparison of ventricular size and function in adult heterozygous *ttn.2* and *ttn.2*/*ttn.1* double mutant zebrafish. | 21 |
| Figure S12 | Normal cardiac morphology and sarcomeric structure in adult heterozygous *ttn.2* zebrafish. | 22 |
| Supplemental Tables | | |
| Table S1 | Primers used for genotyping of mutant zebrafish lines. | 23 |
| Table S2 | Primers used for qPCR analysis. | 24 |
| Table S3 | Primers used for in situ hybridization (ISH) probes. | 26 |
| Table S4 | Antigen sequences for antibody production. | 27 |
| Table S5 | Zebrafish *ttn.2* exons targeted in the mutant lines evaluated and corresponding human *TTN* exons. | 28 |
| Table S6 | Echocardiographic assessment of adult *ttn.2* zebrafish. | 29 |
| References | | 30 |

### Materials and Methods

#### Zebrafish husbandry

Zebrafish (Danio rerio) were raised at a density of 10-15 fish per 3L tank containing fresh system water at 28°C and were maintained as per standard procedures [1]. All experiments using zebrafish were performed in compliance with relevant laws and institutional guidelines in accordance with: (i) protocols approved by the Garvan Institute of Medical Research/St Vincent's Hospital Animal Ethics Committee and the Institutional Biosafety Committee (Australia) and (ii) licenses held under the UK Animals (Scientific Procedures) Act 1986 and later modifications (United Kingdom).

#### Generation of zebrafish lines

Zebrafish lines carrying 6 truncating mutations in different regions of the *ttn.2* gene were generated (Figure S1). Exon numbers are based on [2]. The *ttn.2*$^{e5}$ line (e5; Figures 1A and S1A) was generated on the AB background using ENU mutagenesis, followed by genome sequencing mapping [3-6]. *ttn.2*$^{e25}$ (e25; Figures 1A and S1B), *ttn.2*$^{e129}$ (e129; Figures 1A and S1D) and *ttn.2*$^{e232}$ (e232; Figures 1A and S1F) lines were generated using clustered regularly interspaced short palindromic repeats (CRISPR) - CRISPR associated protein 9 (Cas9) - mediated gene editing. The e25, e129, and e232 lines were maintained on a Tupfel Long Fin (TupLF) background. The *ttn.2*$^{e105}$ line (e105; Figures 1A and S1C) was generated on the TE background using transcription activator-like effector nuclease (TALEN)-mediated genetic engineering. The *ttn.2*$^{e201}$ line (e201; Figures 1A and S1E), has previously been described [7]. The e5, e105, and e201 lines were maintained on a TE background. Variants were confirmed by Sanger sequencing. Sequencing of *ttn.1* at the orthologous locus showed unaltered *ttn.1* sequence in the *ttn.2* mutants.

The *ttn.1* e7 line (Figure S2) was obtained by injecting zygotes with EnGene Spy Cas9 NLS protein (New England Biolabs, Ipswich, Massachusetts, USA) with Alt-R CRISPR-Cas9 crRNA and tracrRNA (Integrated DNA Technologies, Inc., Coralville Iowa, USA) to target GCTTGCGTTGACATGATAGA**AGG** in the Z-disk region of *ttn.1*. The $ttn^{xu071}$ line (kindly provided by Xiaolei Xu) has a double truncation (*ttn.2* A-band and *ttn.1* Z-disk) and has been described in detail [8]. Genotyping of all lines was performed by polymerase chain reaction (PCR) amplification of genomic DNA extracted from whole embryos or adult fin clips, followed by Sanger sequencing or restriction enzyme digest, using the primers and enzymes listed in Table S1.

**qRT-PCR**

Total RNA from pooled wild-type (WT), as well as homozygous and heterozygous *ttn.2* embryos (30 embryos/sample, n=5 samples), or from pooled WT and heterozygous *ttn.2* adult hearts (2 hearts/sample, n=5 samples) was extracted using TRIzol (Sigma-Aldrich, St. Louis, Missouri, USA) and the RNeasy Micro Kit (Qiagen, Hilden, Germany) as described [9]. Purified RNA (1000ng) was used to generate cDNA using the Superscript III First-Strand Synthesis System (Invitrogen, Burlingame, California, USA). Relative qPCR was carried out in 384-well plates using a Light Cycler 480 thermal cycler (Roche, Basel, Switzerland) with primers listed in Table S2. Gene expression was normalized to the expression level of the housekeeper genes Tmem50a and Ube2a using ΔΔCt (cycle threshold) values and graphed relative to transcript expression in WT fish or as absolute expression.

***In situ* mRNA hybridization, and immunofluorescence**

In situ mRNA hybridization was performed as described [10]. Probes used were kinase, N2A, and N2B for *ttn.2* and *ttn.1,* and Cronos [11] (primers listed in Table S3). For immunohistochemistry, primary antibodies were: α-actinin, 1:500 (A7811; Sigma-Aldrich), actin (Actc1a), 1:100 (GeneTex, Irvine, California, USA), myosin binding protein C (MyBP-C), 1:200 [12], an antibody made against human cardiac C1-C2 that recognises cardiac, slow (strongly) and fast (weakly) skeletal isoforms in zebrafish, Myom1 (myomesin B4, 1:100 [13]), Myom2 (M-protein, AA259) [14], sarcomeric myosin heavy chain (MyHC; A4.1025), 1:10 [15], fast myosin light chain (fast MyLC, F310), 1:10 [16], titin (T12), 1:10 [17], titin (Z1Z2), 1:100 [18], Ttn.2 (Ttn.2 M6, see below), 1:1000, Ttn.1 (Ttn.1 M8-M9, see below), 1:2000. Secondary antibodies were mainly Alexa dye-conjugated (Thermo Fisher Scientific, Waltham, Massachusetts, USA), as well as Goat anti IgA FITC (F9384, Sigma-Aldrich) and Cy3 AffiniPure Goat Anti Mouse IgG, Fcγ fragment specific (Jackson ImmunoResearch Laboratories Inc., West Grove, Pennsylvania, USA). Samples for immunohistochemistry were fixed and stained as previously described [19] and imaged on a Zeiss LSM510. Samples were mounted in agarose and photographed on a Zeiss Axiophot with Axiocam (Carl Zeiss, Oberkochen, Germany) using Zen software (Carl Zeiss).

**Antibody production and testing**

To generate specific antibodies to M-band zebrafish Ttn.1 and Ttn.2, we identified unique regions in M-band Ttn.1 and Ttn.2 chosen from sequence alignments (Table S4). The M6 Ttn.2 antibody was designed for a unique region that shares high homology to human M6 but is missing from Ttn.1. The M8-M9 Ttn.1 antibody was designed for a region homologous to the human Mis6-M8-M9-Mis7 region in Ttn.1. The cDNA constructs for protein expression were amplified from embryonic zebrafish cDNA and primer design was based on *ttn.2* and *ttn.1* sequences (GenBank accession no. DQ649453.1). *ttn.2* or *ttn.1* DNA was cloned into a modified pET vector with an N-terminal $His_6$-tag and TEV cleavage site. The identity of the derived constructs was verified by DNA sequencing. Ttn.2 is4-M6 and Ttn.1 is6-M8-M9-is7 fragments were expressed in the *Escherichia coli* strain BL21-CodonPlus (DE3)-RIPL (Agilent Technologies, Santa Clara, California, USA). Protein was expressed in auto-induction media containing lactose to drive expression of the T7 promoter and cells were harvested by centrifugation and lysed using B-PER Bacterial Protein Extraction Reagent (Thermo Fisher Scientific). Protein was purified by nickel-affinity purification and gel filtration chromatography. After TEV protease cleavage of the His6-tag, Ttn.2 is4-M6/Ttn.1 is6-M8-M9-is7 fragment were used for immunization of rabbits and polyclonal sera were collected (Eurogentec, Seraing, Belgium). The antigen was coupled to NHS-activated Sepharose 4 Fast Flow beads (GE Healthcare, Chicago, Illinois, USA) and serum was affinity-purified as previously described [20].

**Titin protein gels & western blots**

Hearts (ventricle and atrium only) from adult wild-type and heterozygous *ttn.2* zebrafish (male, ~14 months old) were snap frozen in liquid nitrogen (n=2 per sample). Hearts were homogenized on ice in a urea buffer as previously described [21]. Samples were loaded onto a 2.0% agarose-stabilized polyacrylamide gel at a concentration of 15-20 μg under denatured and reduced conditions. Following SDS- polyacrylamide gel electrophoresis (PAGE), gels were either stained immediately using Coomassie blue to visualize total protein or transferred onto 0.2 μm PVDF membranes which were then immunostained for titin. Antibodies used: anti-*TTN* mouse monoclonal antibody (2F12, 1:1000, Abnova, Taipei, Taiwan, #H00007273-M07A), anti-mouse HRP (1:15000, Cytiva, Marlborough, Massachusetts, USA, #GENA931).

**Electron microscopy**

Muscle contraction was inhibited using 20 mM 2,3-butanedione monoxime and embryos were fixed in a solution of 4% paraformaldehyde/2.5% glutaraldehyde in PBS for 1 h on ice, postfixed in a 1% osmium tetroxide in PBS solution for 30 m on ice, followed by graded dehydration in ethanol on ice. Sections were stained with uranyl acetate or UranyLess (Electron Microscopy Services, Hatfield, Pennsylvania, USA). Electron microscopy was carried out using a JEOL JM1400 transmission electron microscope in the Centre for Ultrastructural Imaging, King's College London.

**Video-microscopy**

Phenotypic evaluation was performed in anesthetized zebrafish embryos at room temperature using video-microscopy as described [7] with a Leica DM IL inverted microscope (Leica Microsystems, Wetzlar, Germany) and a Nikon DS-Qi1MC camera (Nikon Instruments, Tokyo, Japan) with NIS Elements AR v.3.1 software (Nikon Instruments). Heart rates were obtained from the video images. End-diastolic (EDA) and end-systolic (ESA) chamber areas were derived from short (a) and long axis (b) diameters according to the formula: $A = \pi * ½ a * ½ b$. Fractional area change (FAC) was derived using the formula: FAC = (EDA-ESA)/EDA [7].

**High frequency echocardiography**

Underwater echocardiography was performed in adult zebrafish aged 9-15 months using the Vevo3100® Imaging Station (VisualSonics, Amsterdam, Netherlands) equipped with a high frequency transducer (MS700D) as described [7, 22]. Male fish were used in this study. We have previously reported that there is considerable variability in echocardiographic measurements in female fish due to technical issues, e.g. distortion of the heart's position due to eggs, and varying body weight depending on the number of eggs and gravity status. The latter confounds use of body weight for chamber size normalization) [22]. Two-dimensional (B-Mode) images, color and pulsed-wave Doppler signals were recorded in the long axis view optimized for either ventricular or atrial assessment, respectively. Image analysis was performed using the

VevoLab$^{TM}$ analysis software package version 5.7.0 (VisualSonics) [7, 22] by a single operator who was blinded to genotype. B-Mode images in the long axis view were used to derive ventricular end-diastolic and end-systolic volumes (EDV, ESV), and maximal atrial size (atrial area, AA), indexed to body surface area (BSA). Speckle tracking analysis of ventricular wall motion was performed with the VevoStrainTM analysis software package (VisualSonics) and used to calculate heart rate, ejection fraction (EF), global longitudinal strain (GLS) and global longitudinal strain rate (GLSR). Pulsed-wave Doppler signals measured include: ventricular outflow tract velocity (VOT), E wave (peak velocity of blood inflow across the atrioventricular valve during early diastole), A wave = peak velocity of blood inflow across the atrioventricular valve during atrial systole, and isovolumic relaxation time (IVRT).

**Adrenaline Stress**

10-15 month old heterozygous e5 (e5+/-) and e105 (e105+/-) mutants and WT siblings were subjected to acute adrenaline stress by submersion in 500 µM epinephrine hydrochloride (E4642, Sigma-Aldrich) for 2h [23]. Systolic and diastolic function were measured at baseline and at the conclusion of adrenaline exposure using high-frequency ultrasound.

**Statistical analysis**

Statistical analyses were performed with GraphPad Prism 7.03 (GraphPad Software, Inc., California, USA) unless otherwise specified. Normality testing was performed using the Shapiro-Wilk method. Two-way ANOVA was performed on echocardiography, ECG data and during quantitation of histology sections to identify genotype effects, treatment effects, and genotype-treatment interactions. All p-values were adjusted for multiple comparisons using Tukey's post-hoc correction. Data points in plots represent biological replicates, with bars representing mean ± SD. The absolute difference between two groups is reported in the text as the delta (Δ) of the mean ± SD (Δmean ± SD). Significance level (α) is $p \leq 0.05$ for all studies.

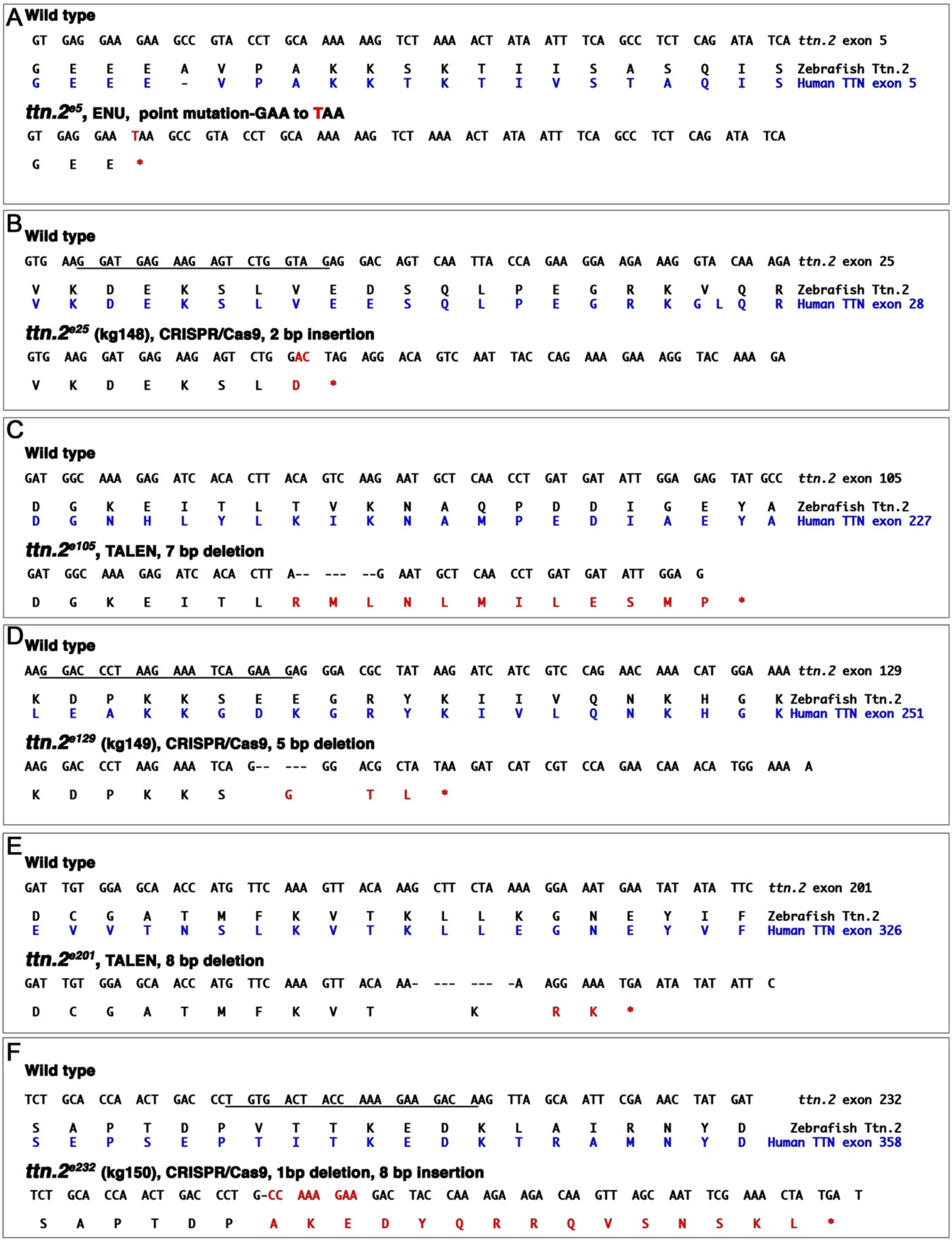


**Figure S1. Generation of zebrafish *ttn.2* truncation variants.** **A** through **F,** Sequence information and methods used to generate *ttn.2* truncation alleles. Rows show DNA and protein sequences for wild-type and *ttn.2* mutant zebrafish (black) and corresponding human protein sequence (blue). CRISPR target sequences are underscored. In the mutant allele sequences, deleted bases (dashed lines) and insertions (red) are shown. Resulting amino acid changes are shown in red. Exon numbering is based on Seeley et al ([2]) for zebrafish and the inferred complete human *TTN* meta-transcript (NM_001267550.2).

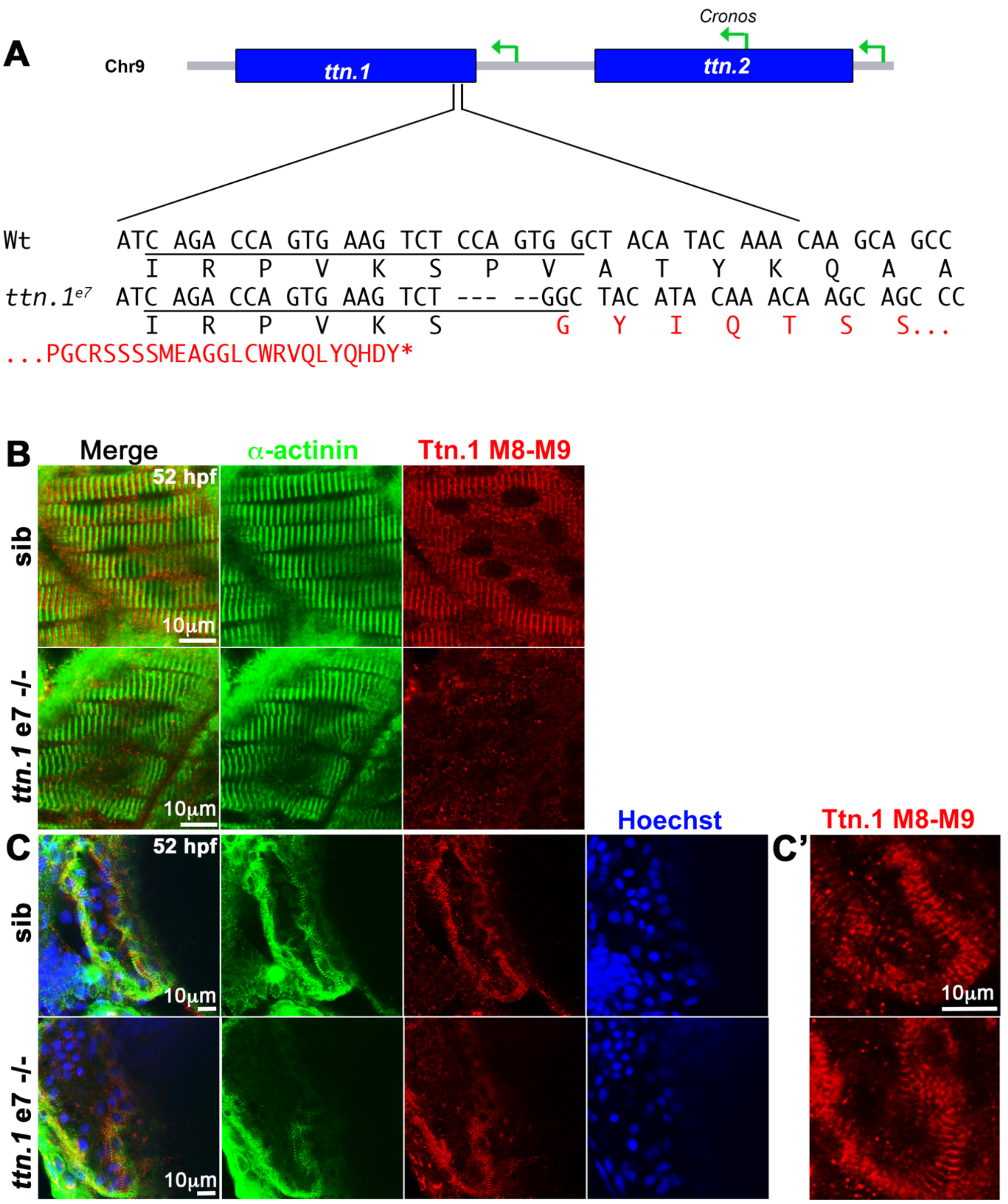


**Figure S2. Zebrafish *ttn.1* truncation variant. A**, DNA and protein sequence for wild-type (WT) and *ttn.1* e7 alleles. CRISPR target sequences are underscored. In the mutant sequences, deleted bases (dashed lines) and altered amino acids (red) are shown. **B** and **C**, Immunofluorescence staining of somitic slow muscle and heart in 52 hpf *ttn.1* e7-/- fish and a genotyped WT sibling for α-actinin and Ttn.1 M8-M9, shown in lateral view anterior to left. Ttn.1 M8-M9 signal is strongly reduced in *ttn.1* e7-/- somitic slow muscle fibers (**B**) but remains strong in cardiomyocytes (**C** and **C'**, higher magnification) indicating that it can also detect Ttn.2.

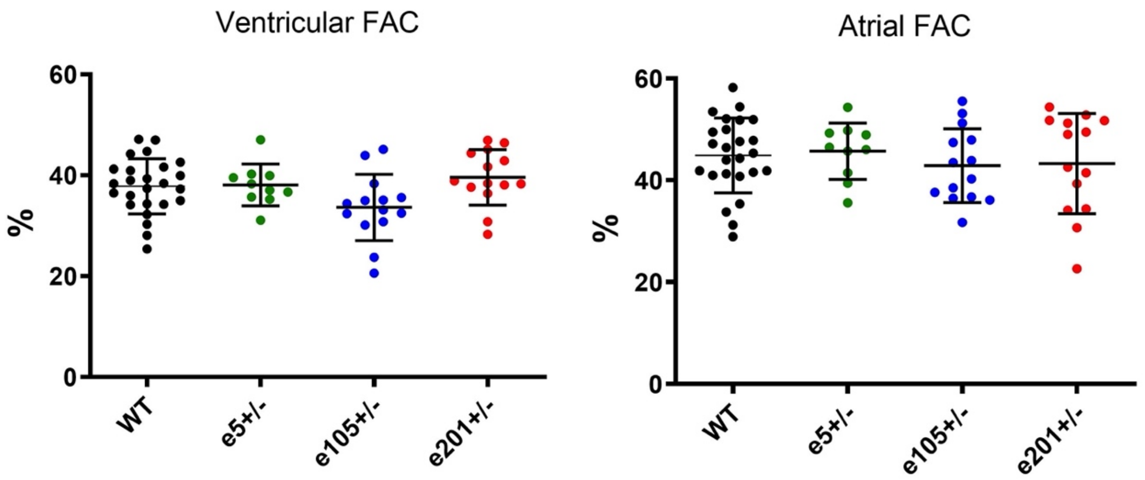


**Figure S3. Cardiac function in heterozygous *ttn.2* embryos.** Ventricular and atrial contractile function were assessed in 3 dpf wild-type (WT) and heterozygous mutant (e5+/-, e105+/-, e201+/-) embryos using video-microscopy. Data are expressed as fractional area change (FAC; %). Unpaired t-tests, mean ± SD.

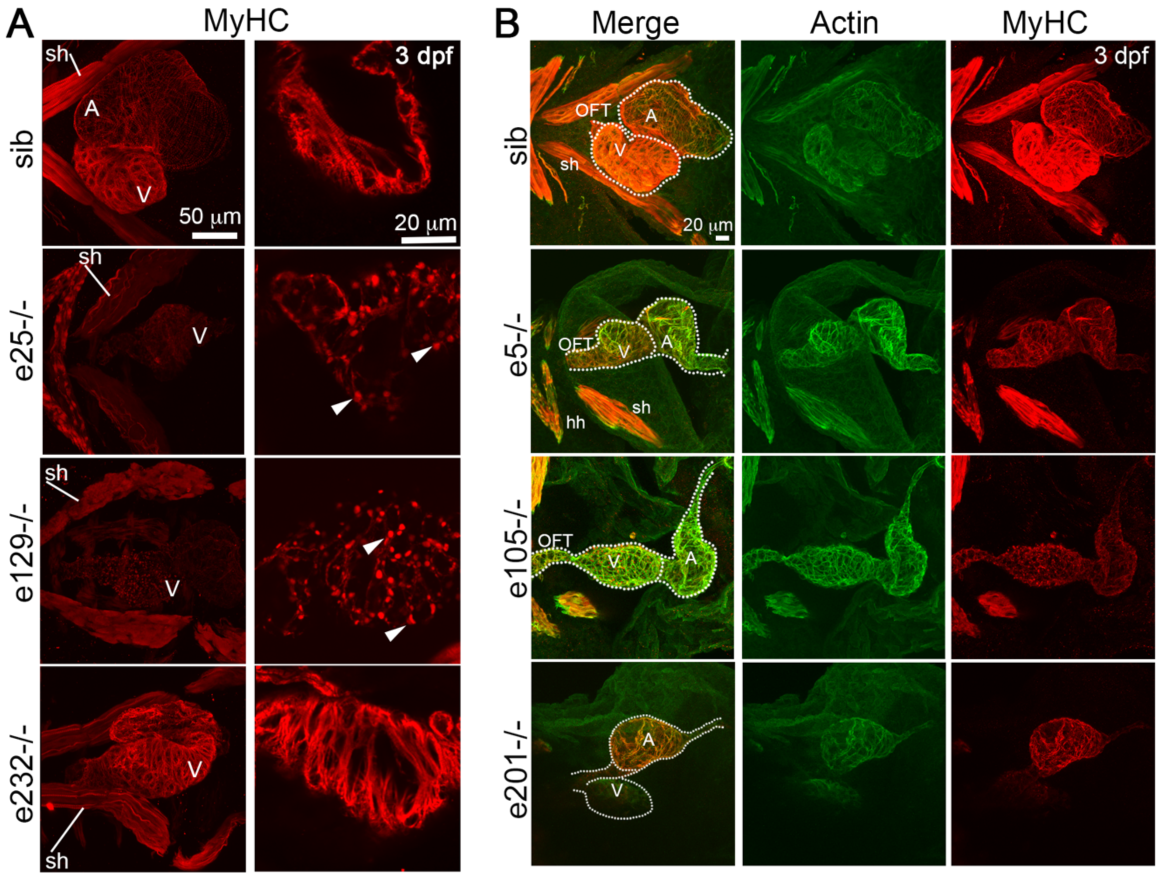


**Figure S4. Abnormal heart development in *ttn.2* mutants. A**, Confocal stacks (left panel) or single scan higher magnification (right panel) of hearts from 3 dpf e25-/-, e129-/- and e232-/- embryos and a sibling (sib) control immunostained for myosin heavy chain (MyHC) shown in ventral view, anterior to left. Heart chambers in e25-/- and e129-/- embryos are small and linear, and the sternohyoideus muscle is pushed outwards by edema. **B,** Confocal stacks of hearts from 3 dpf e5-/-, e105-/- and e201-/- embryos immunostained for actin and MyHC shown in ventral view, anterior to top. Heart chambers in these mutants are also small and linear. A denotes atrium, hh, hyohyoideus, OFT, outflow tract, sh, sternohyoideus, and V, ventricle.

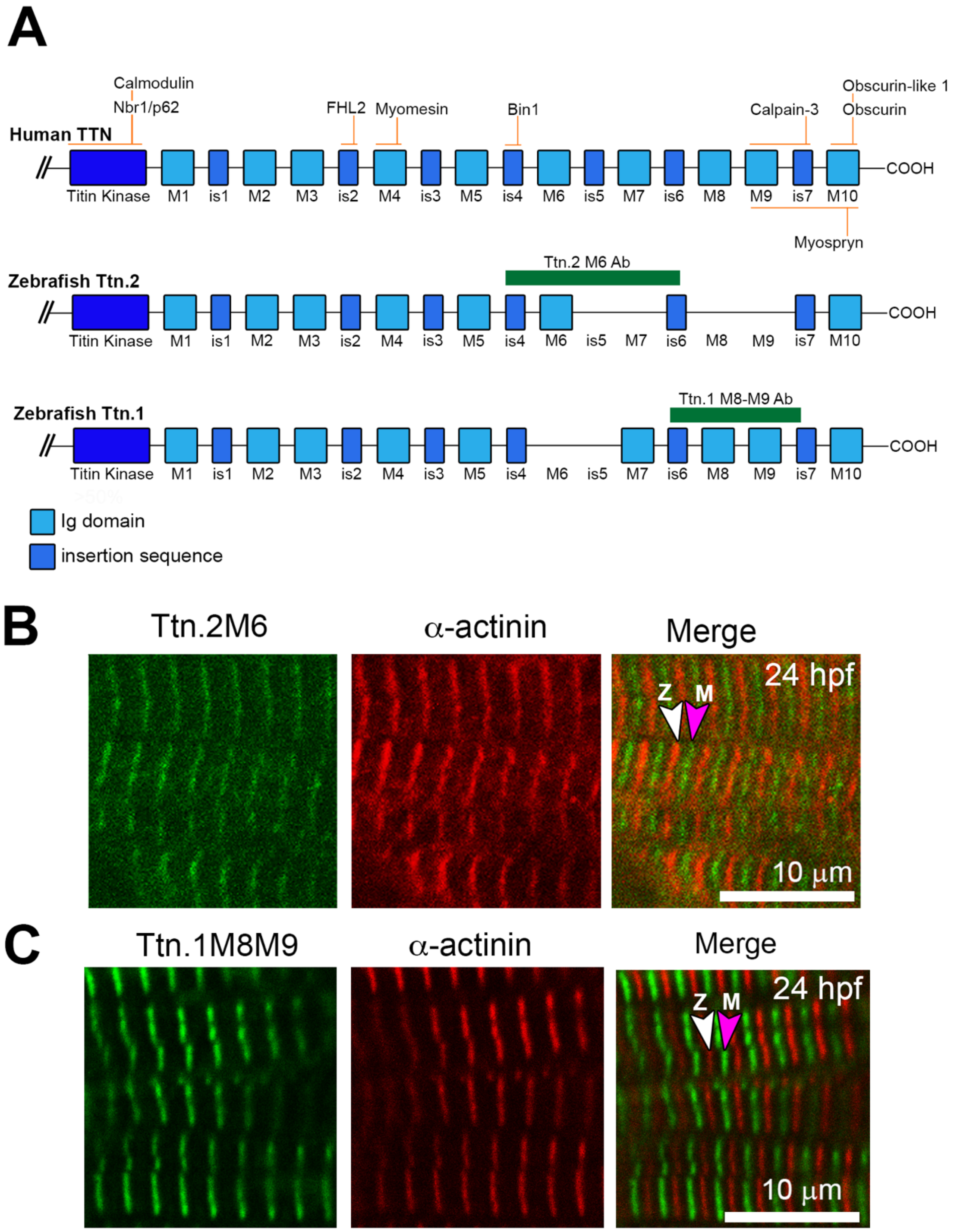


**Figure S5. Differences between the Ttn.2 and Ttn.1 M-band regions and specific antibodies. A**, Comparison of M-band protein domains in human titin and zebrafish Ttn.2 and Ttn.1. Immunoglobulin-like domains M1 to M10 (light blue), unique insertion sequences 1 to 7 (is; dark blue) and antibody epitopes made in this study (green bars) are shown. Known sites of interacting proteins are denoted above. **B** and **C**, Immunofluorescence staining of somites from wild-type embryos at 24 hpf using Ttn.2 M6, Ttn.1 M8-M9, and α-actinin. Both Ttn.2 M6 and Ttn.1 M8-M9 detect Ttn protein localized to the M-band (M) in sarcomeres alternating with α-actinin in the Z-disk (Z).

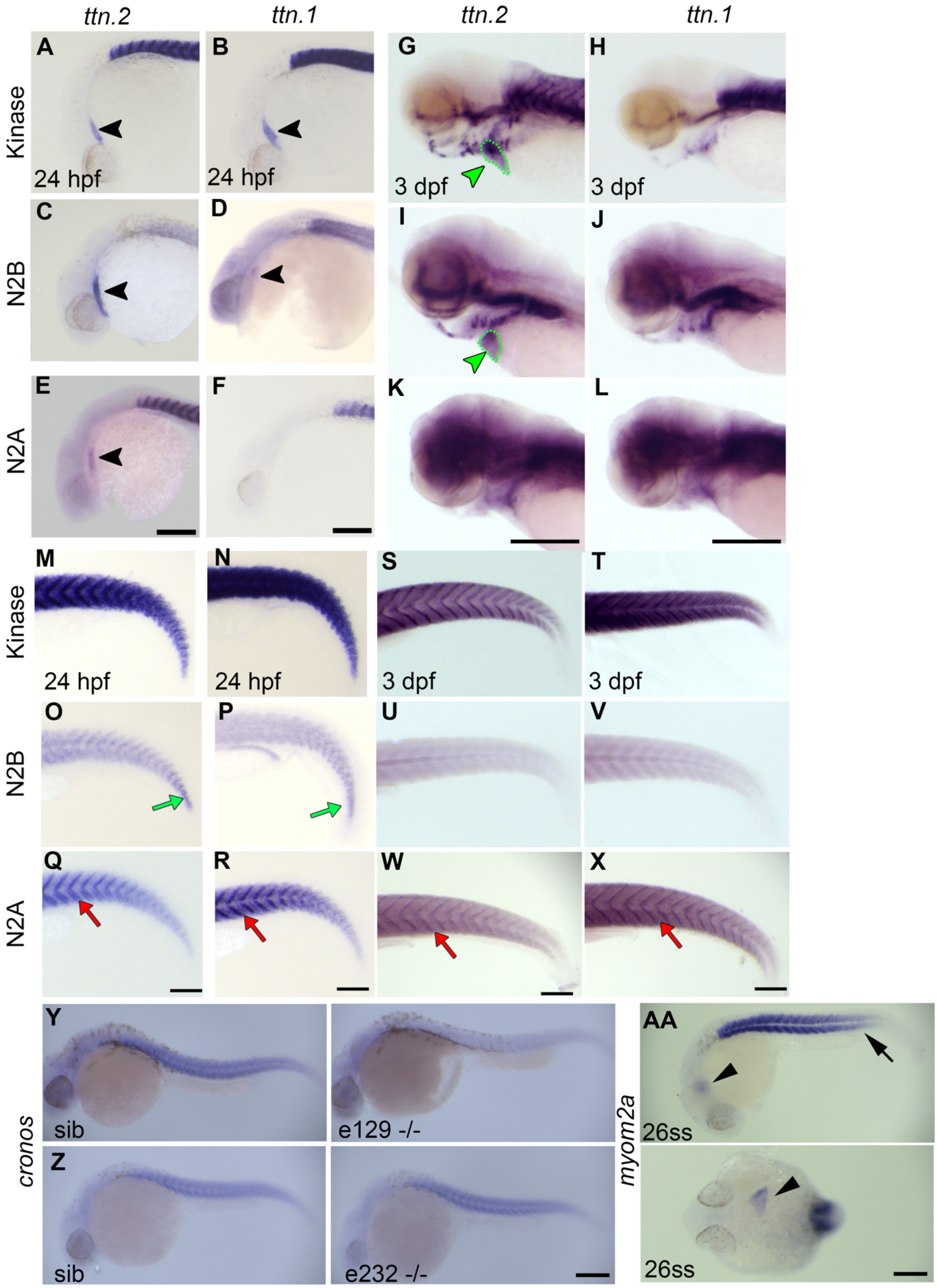
ttn.2
ttn.1
ttn.2
ttn.1
Kinase
N2B
N2A
A
B
G
H
C
D
I
J
E
F
K
L
24 hpf
24 hpf
3 dpf
3 dpf
Kinase
N2B
N2A
M
N
S
T
O
P
U
V
Q
R
W
X
24 hpf
24 hpf
3 dpf
3 dpf
cronos
Y
Z
sib
e129 -/-
sib
e232 -/-
myom2a
AA
26ss
26ss

**Figure S6. Expression of titin in embryonic somitic muscle and heart.** Whole mount in situ hybridization using riboprobes for kinase, N2B and N2A of *ttn.2* and *ttn.1* (purple) in wild-type embryos at 24 hpf (**A-F**, **M-R**) or 3 dpf (**G-L**, **S-X**), lateral view, anterior to left. At 24 hpf, both *ttn.2* and *ttn.1* show expression (black arrowheads) with kinase and the N2B probes (**A,C** and **B,D**, respectively) but not with *ttn.1* N2A (**E,F**). By 3 dpf, expression (green arrowheads) in the heart (green dotted outline) is only detected with *ttn.2* kinase (**G**) and *ttn.2* N2B probes (**I**) but not with *ttn.2* N2A (**K**) or any of the *ttn.1* probes (**H,J,L**). At 24 hpf, *ttn.2* kinase and *ttn.1* kinase are strongly expressed in somites (**M,N**); *ttn.2* and *ttn.1* N2B are diffuse throughout somites (**O,P**) with higher levels in the most caudal somites (green arrows); *ttn.2* and *ttn.1* N2A are diffusely present (**Q,R**) and concentrated along somite borders (red arrows). At 3 dpf, both *ttn.2* and *ttn.1* kinase are present throughout the somites (**S,T**); *ttn.2* N2B and *ttn.1* N2B are only weakly expressed (**U,V**); *ttn.2* and *ttn.1* N2A are also expressed (**W,X)**, with stronger concentration near somite borders (red arrows). **Y** and **Z**, Wholemount *in situ* hybridization in 24 hpf embryos using riboprobe for *cronos* shows a severe reduction of Cronos expression in e129-/- mutants (**Y**) but not in e232-/- mutants (Z) when compared with siblings. **AA.** Wholemount *in situ* hybridization in 22 ss (somite stage) wild type embryos using riboprobe for *myom2a,* lateral view (top panel) and dorsal view (bottom panel), anterior to left. Expression is typical of differentiated fast skeletal muscle and lacking in the tail (arrow) as well as in the developing heart (arrowheads).

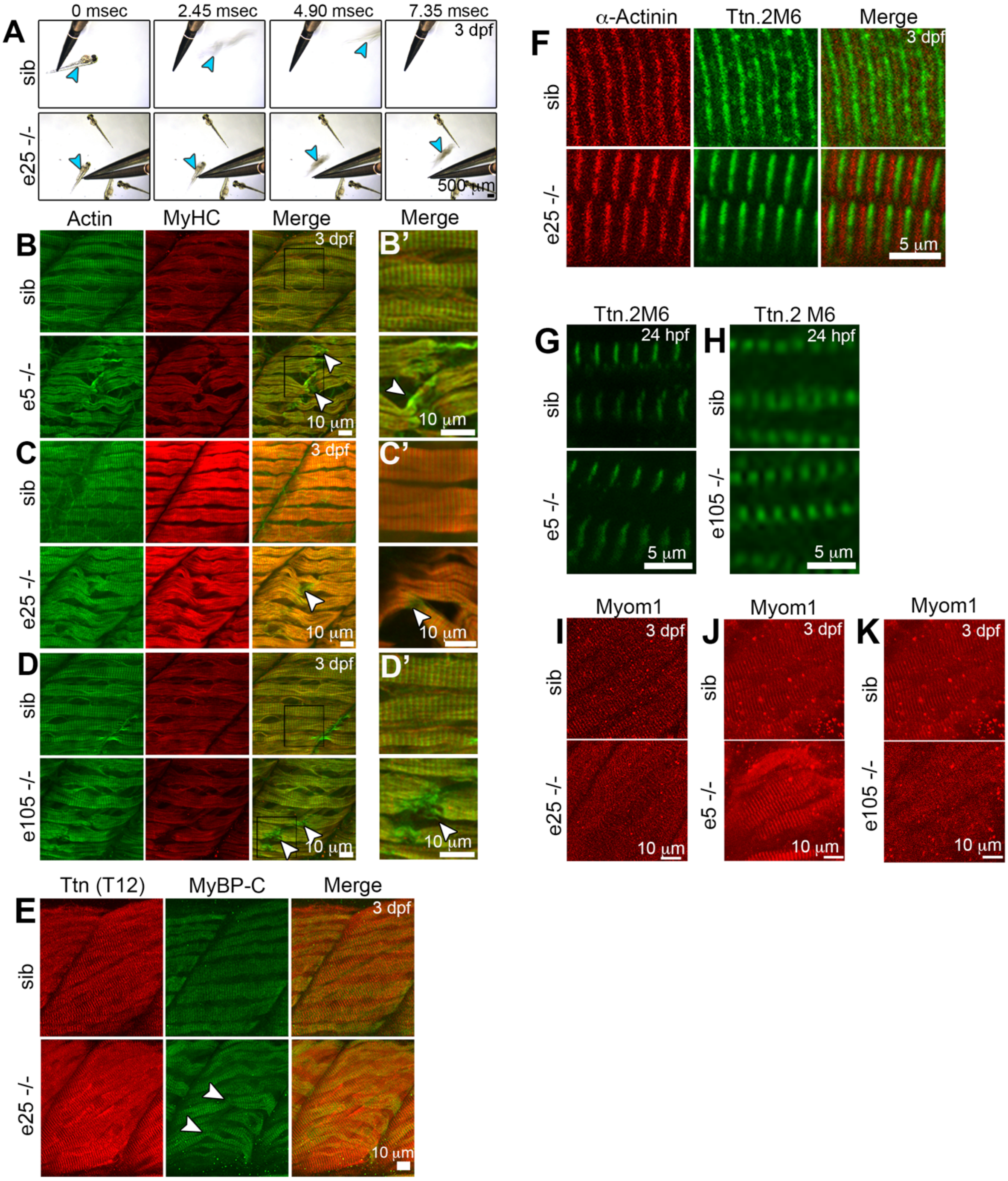

A
0 msec
2.45 msec
4.90 msec
7.35 msec
3 dpf
sib
e25 -/-
500 µm
Actin
MyHC
Merge
Merge
B
B'
e5 -/-
10 µm
C
C'
D
D'
e105 -/-
E
Ttn (T12)
MyBP-C
F
α-Actinin
Ttn.2M6
5 µm
G
H
24 hpf
Ttn.2 M6
I
J
K
Myom1

**Figure S7. Homozygous embryos with *ttn.2* truncation sites proximal to the *cronos* promoter have abnormalities in muscle fiber structure and motility. A**, Sequential brightfield images (2.45 msec apart) of 3 dpf e25-/- and sibling embryos showing response to mechano-stimulation. Each fish (blue arrowhead) is touched with forceps at 0 msec and tracked. In response to touch, e25-/- embryos respond but move more slowly and only a short distance compared with siblings. **B** through **D**, Confocal stacks of somitic slow muscle of 3 dpf e5-/- (**B**), e25-/- (**C**) and e105-/- (**D**) embryos and siblings immunostained for actin and myosin heavy chain (MyHC), ventral view, anterior to top. Mutants show variable fiber breakage and damage (white arrowhead) with greater effects in MyHC than actin. Higher magnification of the black box region (**B',D'**) or different fibers (**C'**) is shown. **E** and **F,** Immunofluorescence staining for Ttn T12 and myosin binding protein C (MyBP-C), α-actinin. and Ttn.2 M6 in 3dpf e25-/- and sibling control embryos, shown in lateral view, anterior to left. Slow sarcomeres in e25-/- embryos show strong MyBP-C and Ttn T12 staining but breaking points in fibers are evident (**E**, white arrowheads). High magnification shows α-actinin and Ttn.2 M6 (likely from Cronos) in e25-/- fibers (**F**). **G** and **H,** Immunofluorescence staining for Ttn.2 M6 in slow muscle fibres of 1 dpf e5-/- (**G**) and e105-/- (**H**) embryos and siblings. Staining patterns are normal in the mutant embryos, presumably detecting Cronos. **I** through **K**, Immunofluorescence staining for myomesin-1 (Myom1) in fast muscle fibers of 3 dpf e25-/- (**G**), e5-/- (**H**) and e105-/- (**I**) embryos and siblings shows normal striation patterns.

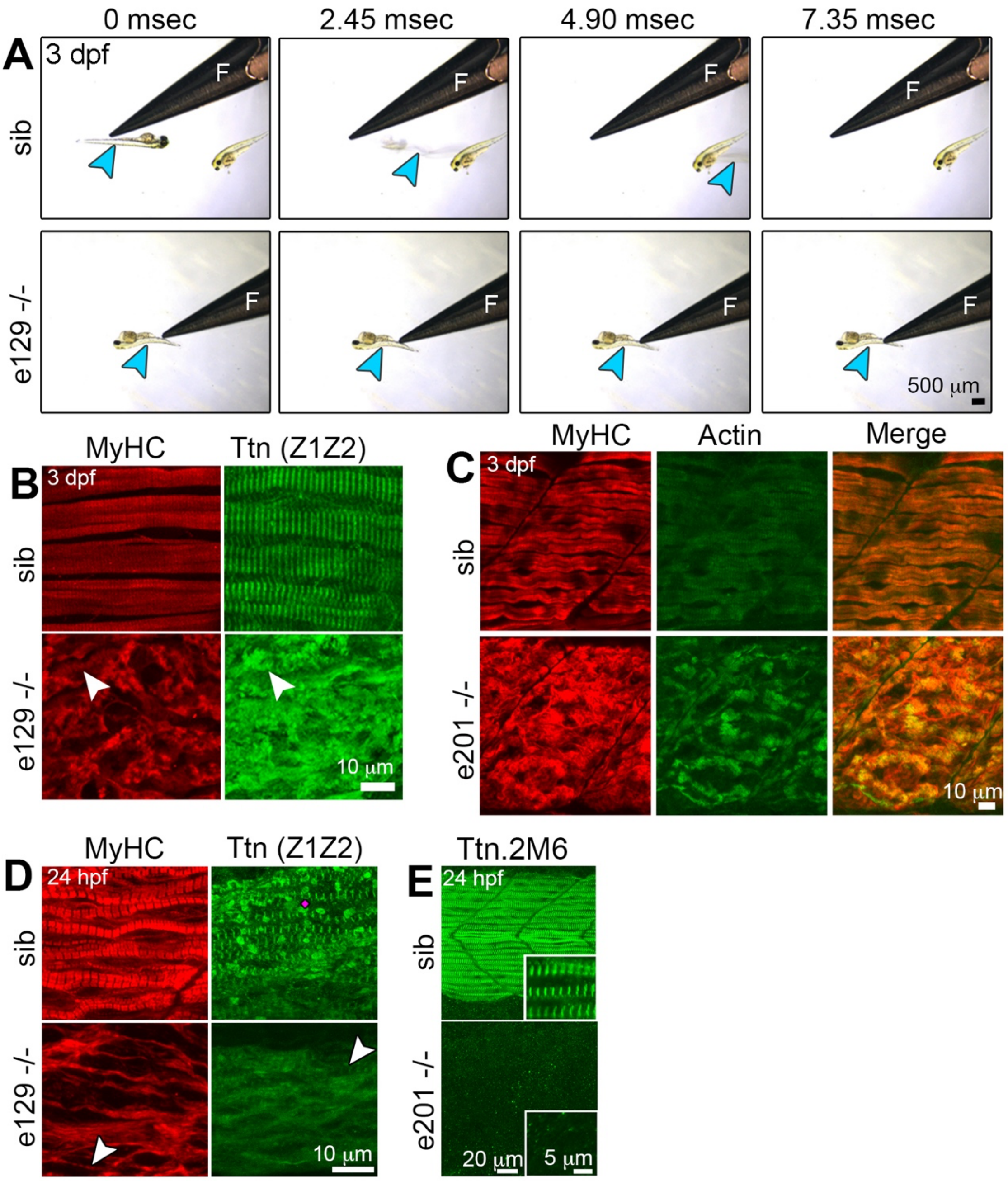


**Figure S8. Homozygous embryos with *ttn.2* truncation sites distal to the *cronos* promoter have severe abnormalities in muscle structure and function. A,** Sequential brightfield images (2.45 msec apart) of 3 dpf e129-/- and sibling (sib) control embryos after mechano-stimulation. Each fish (blue arrowhead) is touched with forceps at 0 msec and tracked. e129-/- mutants are paralyzed and do not respond to touch. **B** through E, Confocal stacks of somitic muscle of e129-/- (**B,D**) and e201-/- (**C,E**) and sibling control embryos at 24 hpf (**D,E**) or 3 dpf (**B,C**) immunostained for titin Z1Z2, myosin heavy chain (MyHC), actin, and Ttn.2 M6, ventral view, anterior to top. MyHC, actin and titin are all disorganized and aggregated in e129-/- and e201-/- embryos (**B,C**). At 24 hpf, fibers in e129-/- embryos are immature and show lack of organized sarcomeres (**D**). No Ttn.2 M6 is detected in e201-/- mutants (**E**).

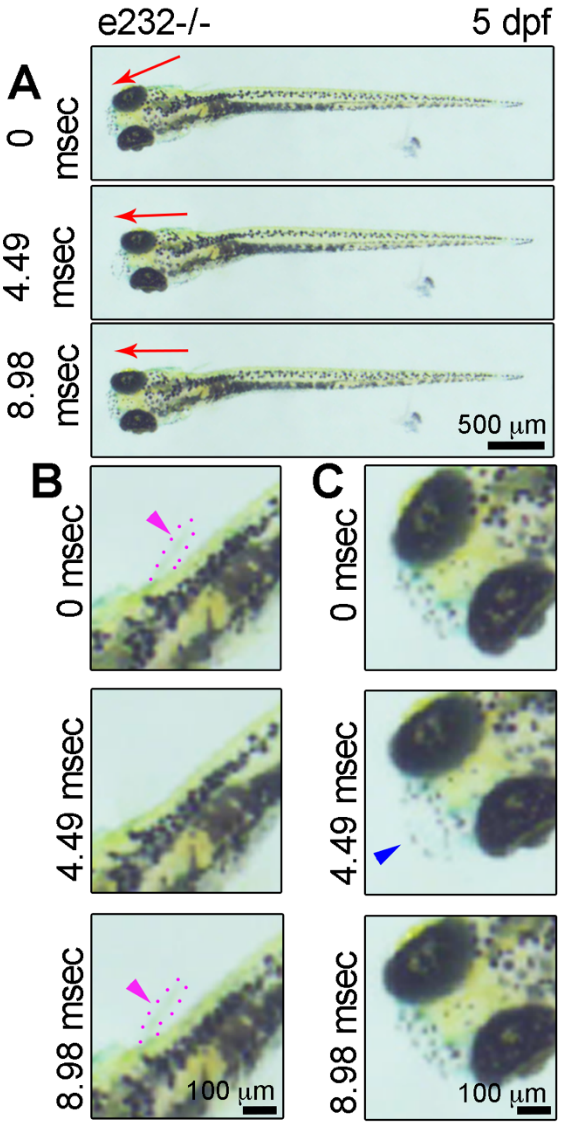


**Figure S9. *ttn.2* e232-/- embryos maintain motility of various muscles.** Sequential brightfield images (4.49 msec apart) of *ttn.2* e232-/- embryos show eye (**A**), jaw (**B**), and fin (**C**) movement at 5 dpf. Red arrows indicate change in eye direction (**A**). Magenta arrowheads (**B**) indicate pectoral fin movement (outlined by magenta dotted line). Opening of jaw is indicated with blue arrowhead (**C**).

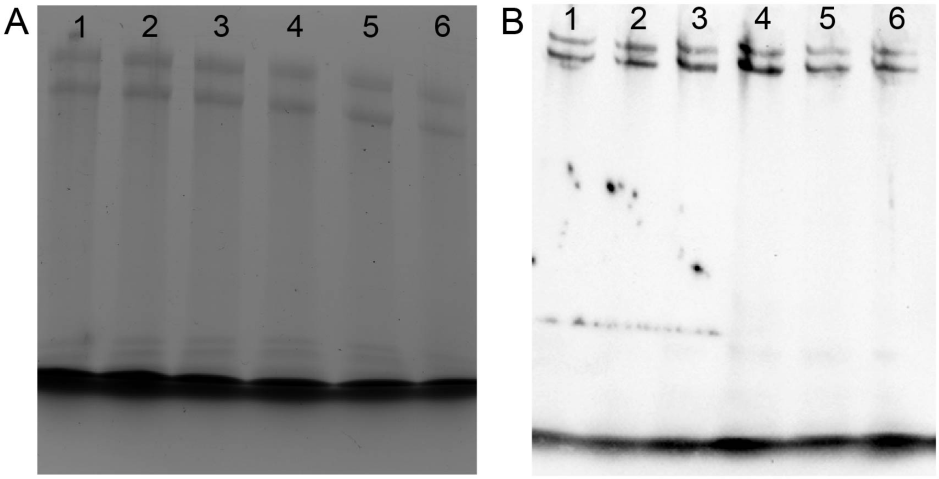


**Figure S10**. **Evaluation of titin levels in adult zebrafish heart.** Cardiac titin protein levels were evaluated using Coomassie staining following SDS-PAGE (**A**) or staining of PVDF membranes using an antibody targeting the Ttn N-terminus (**B**). Band densitometry showed no significant difference in titin protein levels across all 5 lines. There was no clear evidence of truncated titin protein on either Coomassie stained gels or Ttn antibody-stained PVDF membranes. Lanes: (1) wild-type, (2) e5+/-, (3) e25+/-, (4) e105+/-, (5) e129 +/-, (6) e201+/.

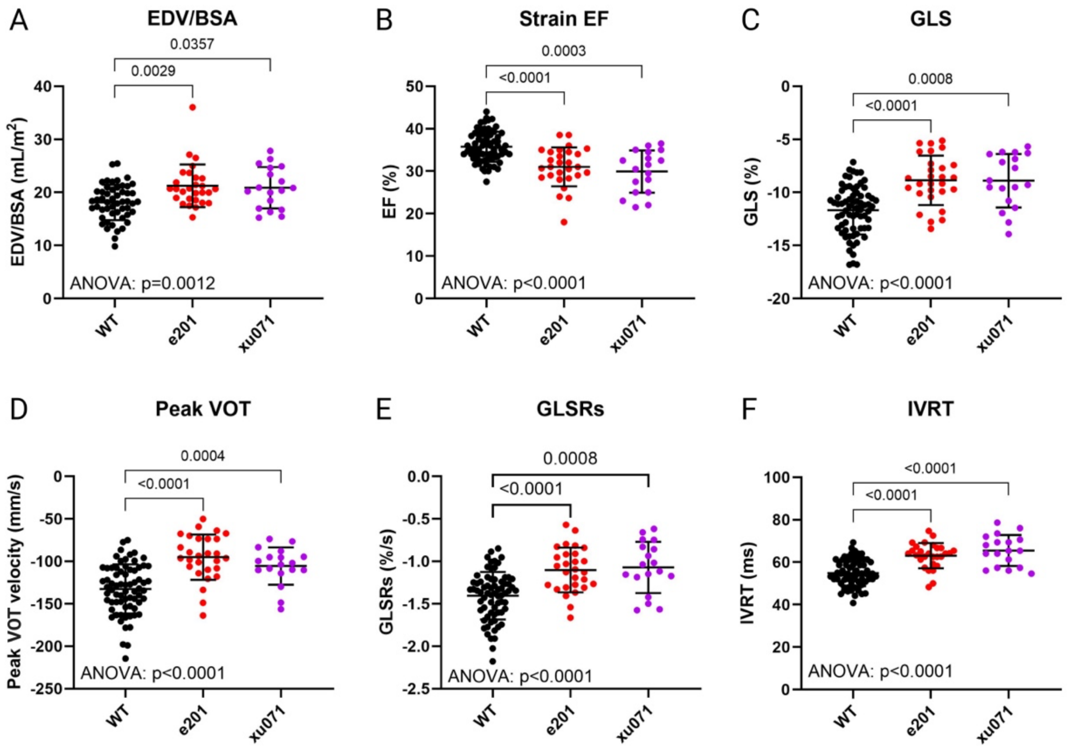


**Figure S11. Comparison of ventricular size and function in adult heterozygous *ttn.2* and *ttn.2/ttn.1* double mutant zebrafish.** Cardiac function was assessed in 9 month-old wild-type (WT), heterozygous single mutant (e201+/-), and heterozygous double mutant (Xu071) fish using high frequency echocardiography. **A,** shows indexed ventricular end-diastolic volume (EDV/BSA); **B**, ejection fraction (EF); **C**, global longitudinal strain (GLS); **D**, peak ventricular outflow tract velocity (VOT); **E**, global longitidudinal strain rate during systole (GLSRs); **F**, isovolumic relaxation time (IVRT). Two-way ANOVA, mean ± SD.

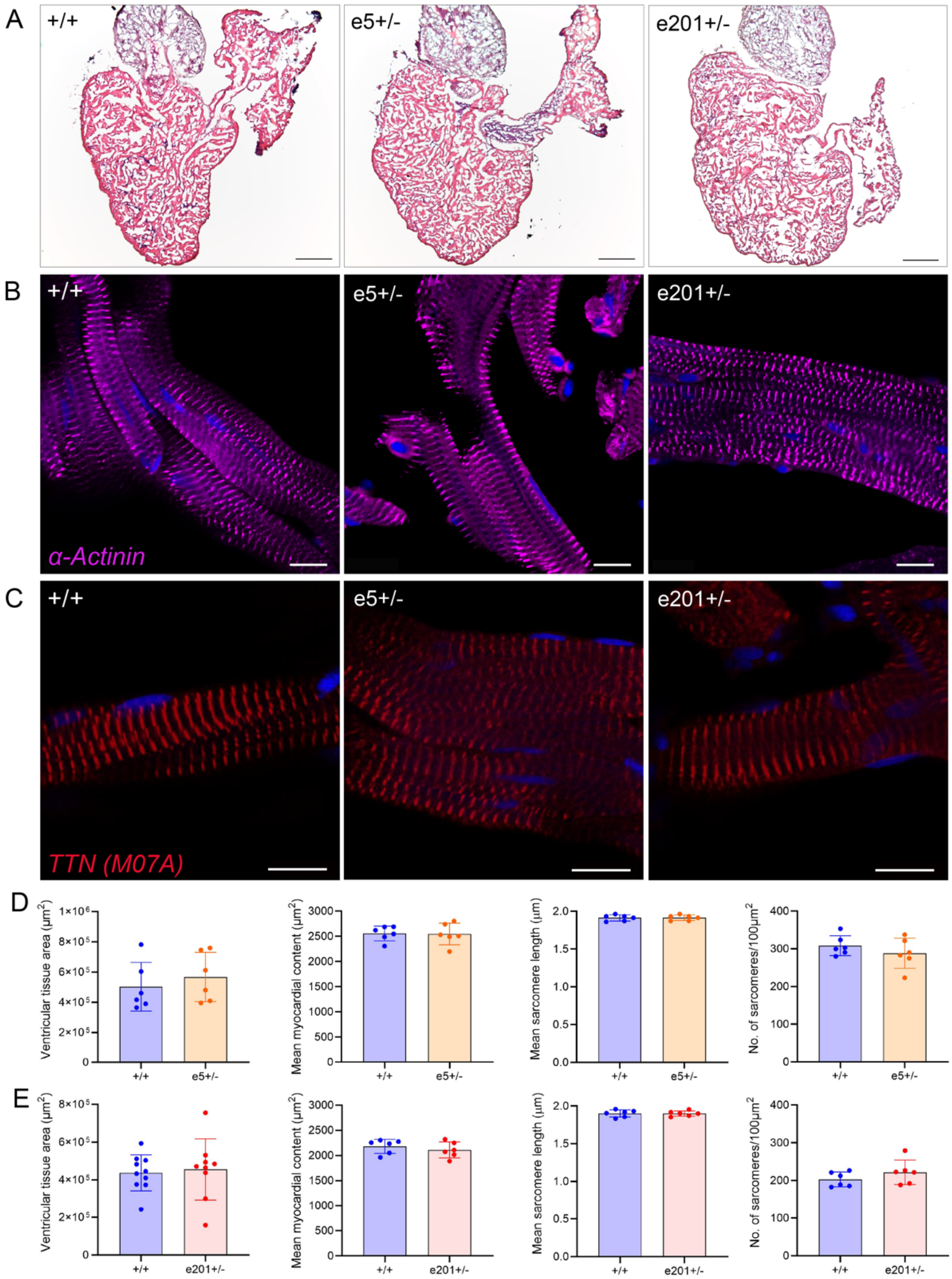


**Figure S12. Normal cardiac morphology and sarcomeric structure in adult heterozygous *ttn.2* zebrafish.** Semi-quantitative analysis of adult heart tissue stained with hematoxylin and eosin (**A**) showed no significant differences in the amount of ventricular trabeculation, assessed by ventricular tissue area or mean myocardial content, in e5+/- (**D**) or e201+/- fish (**E**) relative to wild-type (+/+) siblings. In ventricular tissue sections stained with α-actinin (sarcomere Z-disk marker) (**B**), or an N-terminal Ttn antibody (**C**), there was no evidence of reduced sarcomere number or abnormal periodicity (**D, E**) in the hearts of e5+/- (**D**) or e201+/- (**E**) relative to wild-type siblings. Scale bars: 100 μm. Data were obtained using image segmentation, with values averaged from 5 sections (ventricular tissue area) or 5 regions of interest (mean myocardial content, mean sarcomere length, number of sarcomeres) per heart, n=6 hearts. Unpaired t-tests, mean ± SD.

**Supplementary Table 1.** Primers used for genotyping of mutant zebrafish lines.

| **Allele** | **PCR primer 1** | **PCR primer 2** | **Restriction Enzyme** |
|---|---|---|---|
| *ttn.2* e5 | TTGTTCAAGATGGAGACCTTTACA | GAAAACTTCAGACACGTTACCTG | MboII |
| *ttn.2* e25 (kg148) | TTGTTGAATGGCTCCATGATGGC | CAATACTCTGACATCTCCAGAATC | BfaI |
| *ttn.2* e105 | TGCCTACAGAAATTCTTTATGGTTGT | AGAGAAAACAAAAACAATCTTCTTCAA | TaaI |
| *ttn.2* e129 (kg149) | GAGGCTGAGTGGTTCTACAATGACA | ATCTTGCTGCCGCCATCACT | Eam1104I |
| *ttn.2* e201 | AACCTATGAGGTTGCTTCAG | ACGTAGGTCTGTGACTCTGC | HindIII |
| *ttn.2* e232 (kg150) | CCTGGCAATGATGGTGGAAGTG | GTCCATTACGTGCAAGTTCCTCAGA | NmuCI |
| *ttn.1* e7 (kg184) | GCAGCTCCACGCCATCAGTC | CAAATGACCTTCCCACTGCTGTTC | BpmI |

**Supplementary Table 2.** Primers used for qPCR analysis.

| | **Primer name** | **Sequence** |
|---|---|---|
| *ttn.2* z-disk | *ttna*_e6_F2 | TGCCACAATGCAAATGCAGG |
| | *ttna*_e7_R2 | AGGGGACTTGACAGGTCTGA |
| *ttn.2* I-band | *ttna*_ex103_F | GCAAAGCTTGGCAACAAGGA |
| | *ttna*_ex104_R | TCCTGGATCGTTGCTGATCG |
| *ttn.2* A-band | *ttna*_e154_F | ACAAACTGGCAGAAATGCTC |
| | *ttna*_e154_R | GTCGACTTTCTCGAGGTTCA |
| *ttn.2* M-band | *ttna*_e231-232_F | CAATCTCGACAAGTATGAACC |
| | *ttna*_e231-232_R | CAGTGATGGTACGTGTTTGAC |
| Cronos | *ttna*_Cronos-F2 | GGAAAGCGCTATGTGGATTT |
| | *ttna*_Cronos-R1 | CCTAAGGGTTAGCGTGTGAGA |
| *ttn.1* A-band | *ttnb*_e115_F | TGGGAAATGCCTCTTATTGA |
| | *ttnb*_e115_R | GATGACTTCGCCCAGAACTA |
| Housekeeping gene | *tmem50a* F_ZF | ATTTGTGTTGATTTTGCTGTCG |
| | *tmem50a* R_ZF | TTCTGTTGCAAACTCTGAGGAA |
| Housekeeping gene | *ube2a* F_ZF | TGACTGTTGACCCACCTTACAG |
| | *ube2a* R_ZF | CAAATAAAAGCAAGTAACCCCG |
| WT allele specific primers | e5 FWD | CCGCTGACTTTCAGATTGTT |
| | e5 REV | CGAGTCTGACGAGTTTGTGA |

| | | |
|---|---|---|
| WT allele specific primers | e25 FWD | GTCTGGTAGAGGACAGTCAATTAC |
| | e25 REV | CAATACTCTGACATCTCCAGAATC |
| WT allele specific primers | e105 FWD | AGACATCAAAACACAAGCC |
| | e105 REV | GGTTGAGCATTCTTGACTG |
| WT allele specific primers | e129 FWD | ATCAGAAGAGGGACGCTATA |
| | e129 REV | CTCTGCAAGCCAGTTACAGT |
| WT allele specific primers | e201 FWD | GTTCAAAGTTACAAAGCTTCTAA |
| | e201 REV | AGGTGCAACAAAAGGATTTC |
| WT allele specific primers | e232 FWD | CCTGGCAATGATGGTGGAAGTG |
| | e232 REV | TGGTAGTCACAGGGTCAGTT |

**Supplementary Table 3.** Primers used for in situ hybridization (ISH) probes.

| **ISH probe** | **Primer name** | **Sequence** |
|---|---|---|
| *ttn.2* kinase | *ttna*_kinase_FWD | CCTGGCAATGATGGTGGAAGTG |
| *ttn.2* kinase | T3-*ttna*_kinase_REV | GGATCCATTAACCCTCACTAAAGGGAATGCATTGTCCTCCCTGTCTTCACT |
| *ttn.1* kinase | *ttnb*_kinase_FWD | CTTGGCCGTGGACAGTTTGGTAT |
| *ttn.1* kinase | T3-*ttnb*_kinase_REV | GGATCCATTAACCCTCACTAAAGGGAAGCAGGAGAAGGACGAGGGACTACT |
| *ttn.2* N2A | *ttna*_N2A_FWD | ACACCATCCAAGCAGAAAAGC |
| *ttn.2* N2A | T3-*ttna*_N2A_REV | GGATCCATTAACCCTCACTAAAGGGAATTCCTTCTTCAAAAACTCG |
| *ttn.1* N2A | *ttnb*_N2A_ FWD | ATCAAAGCAGAAAAGTCCTAAG |
| *ttn.1* N2A | T3-*ttnb*_N2A_REV | GGATCCATTAACCCTCACTAAAGGGAATCACCCTTCTCCTTCTTCAG |
| *ttn.2* N2B | *ttna*_N2B_ FWD | AAGCATACAGTGTCAGTGGATTTACAGC |
| *ttn.2* N2B | T3-*ttna*_N2B_REV | GGATCCATTAACCCTCACTAAAGGGAAGTCTGAGGATACTCGCCTTC |
| *ttn.1* N2B | *ttnb*_N2B_ FWD | GGAGGTTATAGTTAGTGGTCC |
| *ttn.1* N2B | T3-*ttnb*_N2B_REV | GGATCCATTAACCCTCACTAAAGGGAAGAGCAAAACAGGCTGCGAGACA |
| *ttn.2 cronos* | *Cronos*_FWD | FGAAAGCGCTATGTGGATTTGTAG |
| *ttn.2* cronos | T7-cronos_REV | GCGTAATACGACTCACTATAGCTTTGGATTCAGGAGAAGACATTG |
| *myom2a* | *myom2a*-FWD | CAGTGCACACCAGAGGGAATCAGA |
| *myom2a* | T7-*myom2a*-REV | TAATACGACTCACTATAGGGAGAGAGGCCTTTGCGATGATTTTTGAC |

**Supplementary Table 4.** Antigen sequences for antibody production.

| Antibody | Antigen amino acid sequence |
|---|---|
| Ttn.2 M6 | SPVPSVKSPEPLVKSPVPSLKSPEPSVKSPVPSVKSPEPQIKSPEPTGIKSPEPRIKSPEG<br>IKSPFRVKSPEPATSLQRVKSPPPLKSPEPTTPQGVKSPIASPPRVKSPPPIKSPEPIASP<br>LRVKSPTGLKSPEPQRAKSPPTVKSPEPIMSPKRMKSPLTVKSPTPSKEAPPKIIQQLKAE<br>AFEDKIRMIFVAESSLREVVWYKDSRKLSQSSHYQIHSSADGTCCLYISDVSEDDQGEYSC<br>EIISEGGAVSRTSFSFVGQVFQAIYTKVTAFVAAHKAVQESVSSKIQGGSEMVI |
| Ttn.1 M8-M9 | AALEGKSELTEEIVKKENTYEEVQSYTEIKASKTQMTISQGQTVTLRASIPEASDVKWILN<br>GAELSNSESYRYGVSGSDHTLTIKSISHHDQGILTCEARTEQGVVKCQFDMTVSATHSGSP<br>SFLVQPHSQNVNEGQNVTFTCEITGEPSPEVEWLKDNAVISITSNMKLSRSKNVYTLEIHN<br>ATIEDSGKFTVKAKNKFGQCSATASLNVLTLVEEPARMIIMEKASDATSMQGSFSAKHVVS<br>KMQESSFSSSSM |

**Supplementary Table 5.** Zebrafish *ttn.2* exons targeted in the mutant lines evaluated and corresponding human *TTN* exons.

| *ttn.2* allele | | Zebrafish | | | Human | | | |
|---|---|---|---|---|---|---|---|---|
| | Variant location | Exon number* | Exon size (bp) | Homology with human exon (%) | Exon number† | Exon size (bp) | Exon PSI‡ (GTeX, %) | Exon PSI (DCM, %) |
| e5 | Z-disk | 5 | 89 | 68.6 | 5 | 86 | 100 | 100 |
| e25 | Proximal I-band | 25 | 1697 | 71.3 | 28 | 1694 | 100 | 100 |
| e105 | Distal I-band | 105 | 276 | 66.7 | 227 | 276 | 98 | 100 |
| e129 | Proximal A-band | 129 | 270 | 64.1 | 251 | 270 | 89 | 100 |
| e201 | Mid A-band | 201 | 17103 | 69.3 | 326 | 17106 | 95 | 100 |
| e232 | Distal A-band | 232 | 5717 | 62.2 | 358 | 5609 | 100 | 100 |

* Zebrafish exon numbering as described in Seeley et al [2] (accession no. DQ649453).
† Human exon numbering in accordance with inferred complete human *TTN* meta-transcript (NM_001267550.2).
‡ Exon percent splice-in (PSI) scores in adult heart tissue in the Genotype-Tissue Expression (GTeX) database and in patients with dilated cardiomyopathy (DCM) [24].

**Supplementary Table 6.** Echocardiographic parameters in adult (12 month-old) *ttn.2* zebrafish.

| **Parameter** | +/+ (n=68) | **e5+/-** (n=30) | *p-value* | **e25+/-** (n=15) | *p-value* | **e105+/-** (n=17) | *p-value* | **e129+/-** (n=15) | *p-value* | **e201+/-** (n=15) | *p-value* | *ANOVA p-value* |
|---|---|---|---|---|---|---|---|---|---|---|---|---|
| Body weight (g) | 0.67 ± 0.08 | 0.65 ± 0.08 | 0.96 | 0.66 ± 0.05 | 1.00 | 0.69 ± 0.08 | 0.98 | 0.66 ± 0.09 | 1.00 | 0.67 ± 0.10 | 1.00 | 0.51 |
| Heart rate (bpm) | 136 ± 19 | 133 ± 17 | 0.99 | 144 ± 17 | 0.86 | 136 ± 15 | 1.00 | 122 ± 17 | 0.16 | 136 ± 14 | 1.00 | 0.028 |
| EDV/BSA ($mL/m^2$) | 18.7 ± 3.3 | 18.5 ± 4.6 | 1.00 | 20.8 ± 4.5 | 0.72 | 18.6 ± 3.1 | 1.00 | 21.3 ± 5.5 | 0.69 | 22.6 ± 4.1 | 0.036 | 0.009 |
| ESV/BSA ($mL/m^2$) | 11.7 ± 2.4 | 11.4 ± 2.2 | 1.00 | 12.8 ± 2.6 | 0.90 | 13.2 ± 2.6 | 0.43 | 13.6 ± 3.4 | 0.52 | 16.5 ± 3.9 | 0.005 | <0.0001 |
| EF (%) | 36.0 ±3.8 | 35.4 ± 4.4 | 1.00 | 31.7 ± 5.2 | 0.10 | 32.6 ± 4.9 | 0.20 | 31.3 ± 4.8 | 0.028 | 29.1 ± 6.2 | 0.010 | <0.0001 |
| GLS (%) | -12.8 ± 2.7 | -13.1 ± 2.5 | 1.00 | -10.8 ± 2.5 | 0.14 | -12.0 ± 2.6 | 0.98 | -10.2 ± 2.5 | 0.023 | -9.2 ± 3.0 | 0.007 | <0.0001 |
| GLSRs (%/s) | -1.37 ± 0.27 | -1.45 ± 0.38 | 0.99 | -1.48 ± 0.37 | 0.98 | -1.31 ± 0.34 | 1.00 | -0.99 ±0.19 | <0.0001 | -1.10 ± 0.19 | 0.001 | <0.0001 |
| Peak VOT (mm/s) | -121.3 ± 24.7 | -112.1 ± 22.0 | 0.63 | -131.8 ± 35.8 | 0.99 | -116.7 ± 24.1 | 0.99 | -89.7 ± 29.5 | 0.015 | -97.6 ± 24.1 | 0.035 | 0.0002 |
| VOT-VTI (mm) | 12.95 ± 4.40 | 13.06 ± 3.07 | 1.00 | 11.04 ± 3.13 | 0.55 | 13.27 ± 4.21 | 1.00 | 9.36 ± 3.64 | 0.039 | 11.07 ± 2.43 | 0.32 | 0.0027 |
| GLSRa (%/s) | 1.42 ± 0.50 | 1.45 ± 0.54 | 1.00 | 1.09 ± 0.33 | 0.046 | 1.25 ± 0.38 | 0.85 | 0.99 ± 0.34 | 0.005 | 1.10 ± 0.31 | 0.042 | <0.0001 |
| Atrial area/ BSA($cm^2/m^2$) | 6.7 ± 1.5 | 6.8 ± 1.9 | 1.00 | 6.2 ± 1.7 | 0.99 | 7.2 ± 1.0 | 0.80 | 8.0 ± 1.7 | 0.18 | 7.9 ± 1.0 | 0.010 | 0.0041 |
| E (mm/s) | 44 ± 15 | 38 ± 11 | 0.37 | 48 ± 13 | 0.98 | 38 ±16 | 0.96 | 37 ± 15 | 0.80 | 26 ± 8 | <0.0001 | <0.0001 |
| A (mm/s) | 294 ± 57 | 280 ± 47 | 0.98 | 287 ± 47 | 1.00 | 266 ± 49 | 0.52 | 282 ± 50 | 0.99 | 232 ± 32 | <0.0001 | 0.0008 |
| E/A | 0.15 ± 0.06 | 0.13 ± 0.04 | 0.55 | 0.17 ± 0.04 | 0.97 | 0.14 ± 0.05 | 1.00 | 0.13 ± 0.05 | 0.88 | 0.11 ± 0.04 | 0.009 | 0.0057 |
| IVRT | 54 ± 8 | 58 ± 11 | 0.80 | 56 ± 6 | 0.98 | 68 ± 8 | <0.0001 | 65 ± 10 | 0.017 | 70 ± 12 | 0.004 | <0.0001 |

Groups compared using non-parametric one-way ANOVA with p-values corrected for multiple comparisons using Dunnett's T3 correction. All other P-values refer to differences between each *ttn.2* line to wild-types (+/+). Data presented as mean ± SD. A, denotes peak velocity of blood inflow across atrioventricular valve during late diastole; bpm, beats per minute; E, peak velocity of blood inflow across atrioventricular valve during early diastole; EDV/BSA, ventricular end-diastolic volume indexed to body surface area; EF, ejection fraction; ESV, ventricular end-systolic volume; GLS, global longitudinal strain; GLSRa, global longitudinal strain rate during atrial contraction; GLSRs, peak systolic global longitudinal strain; IVRT, isovolumic relaxation time; VOT, peak velocity of blood outflow from across bulbo-ventricular valve; VOT-VTI, ventricular outflow tract-velocity time integral.
